# Bridging *in vitro* and *in vivo* insights: A PKPD model for effective phage therapy against multidrug-resistant *Pseudomonas aeruginosa*

**DOI:** 10.1101/2024.11.30.626102

**Authors:** Jun Seok Cha, Kyungnam Kim, Hwa Jeong You, Dasom Kim, Hyun Hee Park, SuJin Heo, Choon Ok Kim, Byung Hak Jin, Dongeun Yong, Dongwoo Chae

## Abstract

Bacteriophage therapy provides a promising solution for multidrug-resistant *Pseudomonas aeruginosa*. However, optimizing phage combinations and dosages remains challenging. Here, we developed and validated a pharmacokinetic-pharmacodynamic (PKPD) model integrating *in vitro* and *in vivo* efficacy data using three phage strains (MP-A, PP-A, and PP-B), administered both alone and in combination. *In vitro* phage resistance was substantially delayed by combining phages that exhibit collateral sensitivity. However, multiple phage resistance eventually emerged. *In vivo,* growth of phage-resistant mutants was minimal. Our model suggests that this outcome was due to an initial phage-induced reduction in bacterial load, followed by immune clearance of the phage-resistant bacteria. Simulations indicated that higher phage exposure at the site of infection caused a faster decline in bacterial levels below the immune clearance threshold and prevented excessive immune activation. Overall, this study offers crucial guidance for treatment strategies and highlights the importance of mathematical modeling in advancing phage therapy.

## INTRODUCTION

Multidrug-resistant (MDR) *Pseudomonas aeruginosa* is recognized as a “priority1: critical” pathogen by the World Health Organization.^1^ In the USA, an estimated 32,600 cases of MDR *P. aeruginosa* infections were reported in 2017, prompting its designation as a “serious threat” in the 2019 report by the Centers for Disease Control and Prevention (CDC).^2^ The European Antimicrobial Resistance Surveillance Network (EARS-Net) reported that 12.6% of samples exhibited resistance to multiple antimicrobial groups.^3^

In response to this growing threat, clinicians have revisited older antibiotics, such as polymyxins, and adjusted dosing strategies for meropenem and aminoglycosides. Recent additions to the antibiotic arsenal include ceftolozane-tazobactam, ceftazidime-avibactam, imipenem-relebactam, and cefiderocol.^4–9^ However, resistance against these newer drugs is already emerging.^10,11^

Bacteriophage therapy offers a promising alternative, with anecdotal successes and support from animal model studies.^12–15^ However, it still lacks definitive clinical evidence and broad acceptance. A key challenge in phage therapy is the rapid development of phage resistance due to genetic mutations and phenotypic changes.^16,17^ To combat this resistance, combination phage therapy, often referred to as “cocktail therapy,” is essential. Nevertheless, not all phage combinations are effective, highlighting the need for systematic strategies to formulate compatible phages.

Another critical factor for successful therapy is ensuring adequate phage exposure. Unlike antibiotics, phage concentration is dynamically influenced by the ability of the phage to replicate within the bacterial population.^18^ Consequently, efficient replication at lower doses can lead to greater phage exposure than higher doses followed by low levels of replication. Simplified models suggest that even low initial phage doses can effectively control target bacterial populations.^19^ However, studies have shown mixed results regarding the optimal phage dose, with some indicating minimal benefit ^20^ or increased resistance at higher doses ^21^ and others reporting better outcomes with higher doses ^22^ ^23^.

Given this conflicting evidence, a deeper mechanistic understanding of phage pharmacokinetics pharmacodynamics (PKPD) is essential. To this end, in the current study, we aimed to develop and validate a robust PKPD model of phage therapy against *P. aeruginosa* using phage strains administered individually and in combinations across varying doses. Our goal was to identify optimal phage cocktail formulations and dosing strategies that enhance clinical outcomes.

## RESULTS

### Bacteriophage characterization

#### Phage identification and structural characterization

MP-A, PP-A, and PP-B, isolated from sewage samples, were collected by the South Korean biotech company Microbiotix for phage therapy applications. Transmission electron microscopy revealed that all three phages belonged to the order *Caudovirales* (tailed bacteriophages).

Specifically, MP-A is classified within the *Myoviridae* family, featuring long, contractile tails, whereas PP-A and PP-B are members of the *Podoviridae* family, characterized by short, non- contractile tails (Figure S1A).

#### Target specificity, lytic spectrum, and genomics

The primary targets for MP-A and PP-B are bacterial lipopolysaccharides (LPSs), whereas PP-A targets bacterial pili. The lytic activities of the phages were tested against the domestic YMC strain of *P. aeruginosa*; results revealed lytic spectra of 60%, 56%, and 54% for MP-A, PP-A, and PP-B, respectively (Figure S1B). Bioinformatics analysis determined genome sizes of 40,000–70,000 bp and gene counts of 50–100 (Figure S1C).

#### Adsorption rate and one-step growth

Adsorption tests using P. aeruginosa 15-4 revealed that all three phages achieved over 90% adsorption within 10 minutes, with a multiplicity of infection (MOI) of 0.001 for MP-A and PP-A and 1.0 for PP-B (Figure S1D). MP-A exhibited the highest adsorption efficiency, with less than 0.1% remaining unadsorbed after 30 minutes, followed by PP-B with less than 1% unadsorbed and PP-A with less than 10% unadsorbed. One-step growth assays revealed the rapid lytic action of PP-A, initiating host cell lysis in under 10 minutes, whereas MP-A and PP-B had lysis times of approximately 40 minutes and 100 minutes, respectively (Figure S1E). Burst size measurements showed that MP-A and PP-A had similar values of approximately 20 and 50, respectively, whereas PP-B exhibited a considerably larger burst size of approximately 200.

### *In vitro* dynamics of mono- and cocktail phage treatments

#### Time-kill assays

We conducted 24-hour *in vitro* time-kill assays against P. aeruginosa 15-4 to examine the kinetics of bacterial growth suppression by phages MP-A, PP-A, and PP-B, both individually and in combinations (Figure S2). These assays showed a dose-dependent suppression of bacterial growth within the initial 10 hours. However, most treatments led to the emergence of resistant bacteria beyond this period. Notably, higher phage doses resulted in enhanced growth of resistant strains during MP-A monotherapy and, to a lesser extent, during PP-A, MP-A + PP-B, and PP-A + PP-B treatments.

The combination of MP-A and PP-A significantly delayed the development of resistance compared to either phage used alone. However, the MP-A + PP-B and PP-A + PP-B combinations were no more effective than their respective monotherapies. A cocktail of all three phages (MP-A, PP-A, and PP-B) showed efficacy comparable to that of the MP-A + PP-A combination, with no additional benefit from including PP-B. In all the tested cocktails, the constituent phages were mixed in 1:1, 3:1, or 1:3 ratios.

### Mathematical modeling of *in vitro* phage-bacteria interactions

#### Model development

We employed mathematical modeling to interpret the *in vitro* time-kill data from phage treatments involving MP-A, PP-A, and PP-B, both individually and in combination. Using a nonlinear mixed-effects framework, we evaluated various models and selected the best-fit model based on statistical criteria and physiological relevance.

The chosen model divided the bacterial population into two subpopulations: phage-susceptible wild-type (W) and phage-resistant mutant (M). The model also assumed that bacteria can shift between proliferating (P) and dormant (D) states, with the latter state being impervious to phage infection (Figure 1A). Following the approach of Nielsen et al., we posited that the transition of proliferating bacteria to the dormant state increased with total bacterial density.^24^ The equations governing these dynamics are detailed in the STAR Methods.

**Figure 1.**
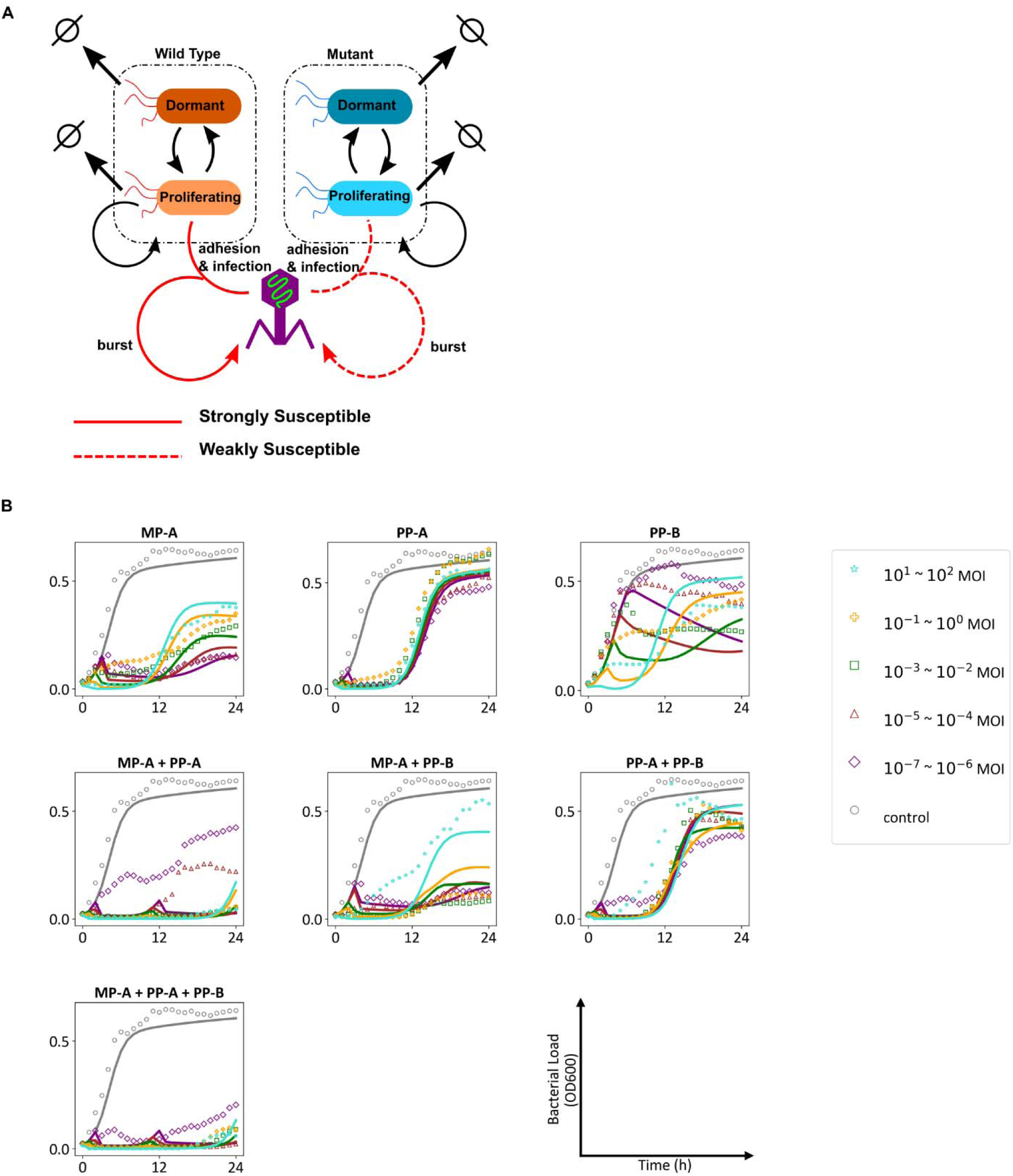
Population dynamics modeling and *in silico* predictions. (A) Model schematic diagram. (B) Typical model predictions overlaid on average observations categorized by dosage cohorts.

#### Model predictions and parameter estimates

The model closely aligned with experimental results, as depicted in Figure 1B. It provided estimates for key parameters, including the bacterial growth rate (k_growth_), natural death rate (k_death_), proliferative capacity (ta9_10_c), rate of transition from the dormant to proliferative state (k_DP_), and the per bacterium phage infection rates (k_inf_) against both W and M types. Moreover, it quantified burst sizes (b) for each phage and estimated the fractions of bacteria resistant to single, double, and triple phage combinations (detailed in Table 1) and included random inter-experimental variability (Table S1).

**Table 1.**
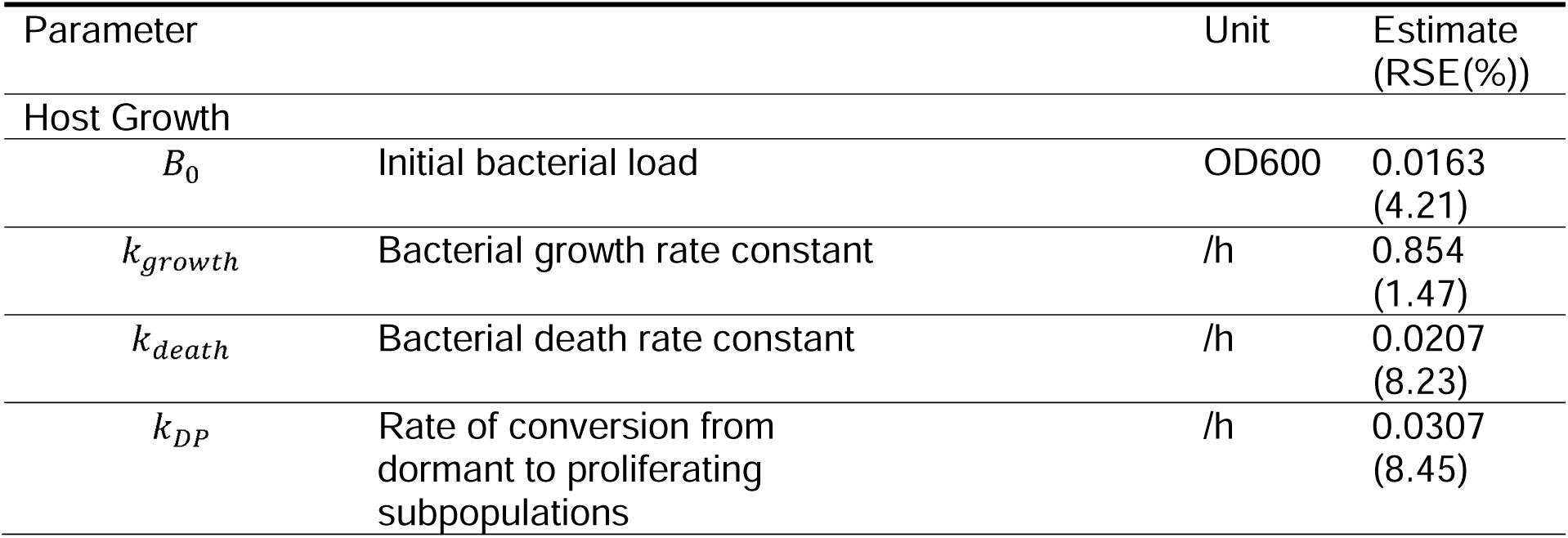
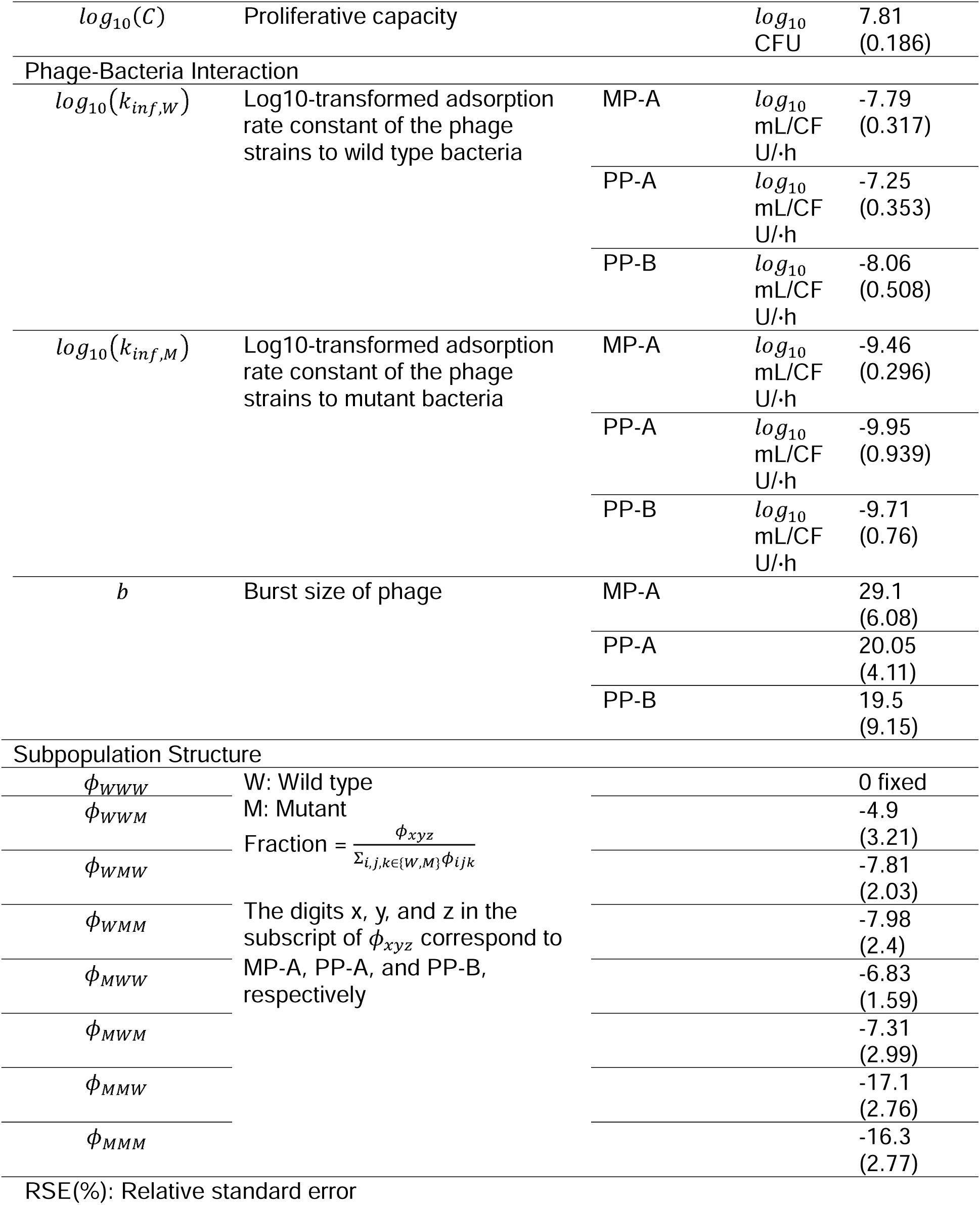
Parameter estimates of the population dynamics model.

The infection rates against W bacteria were the highest for PP-A (10^-7.25^ mL/CFU/h), followed by MP-A (10^-7.79^ mL/CFU/h) and PP-B (10^-8.06^ mL/CFU/h). The infection rates against M bacteria were 1 to 3 logs lower than those against W variants, suggesting that considerably higher phage concentrations are required to effectively target them. Among the phages, MP-A exhibited the highest infection rate against M variants (10^-9.46^ mL/CFU/h), followed by PP-B (10^-9.71^ mL/CFU/h) and PP-A (10^-9.95^ mL/CFU/h).

According to our model estimates, PP-A had the lowest probability of resistance (7.39 x 10^-4^), followed by MP-A (1.73 x 10^-3^) and PP-B (8.38 x 10^-3^) (Table 2). Under the null hypothesis of independent resistance acquisition, the expected double-resistance probabilities for MP-A + PP-A, MP-A + PP-B, and PP-A + PP-B were 1.28 x 10^-6^, 1.45 x 10^-5^, and 6.19 x 10^-6^, respectively. However, the actual estimated probabilities of double resistance for these combinations were 1.18 x 10^-7^, 6.63 x 10^-4^, and 3.38 x 10^-4^, respectively, demonstrating significant deviations from the expected values. These deviations indicated that MP-A + PP-B and PP-A + PP-B exhibited cross-resistance, whereas MP-A + PP-A demonstrated collateral sensitivity, shedding light on the unique synergy between MP-A and PP-A. This comparison allowed for an objective and quantitative assessment of cocktail compatibility.

**Table 2.** Cross-resistance probability.

| Resistance | Estimated Value | Expected Value Under Independent Mutations |
| --- | --- | --- |
| MP-A | $1.73 \times 10^{-3}$ | - |
| PP-A | $7.39 \times 10^{-4}$ | - |
| PP-B | $8.38 \times 10^{-3}$ | - |
| MP-A +<br>PP-A | $1.18 \times 10^{-7}$ | $1.28 \times 10^{-6}$<br>$((1.73 \times 10^{-3}) \times (7.39 \times 10^{-4}))$ |
| MP-A +<br>PP-B | $6.63 \times 10^{-4}$ | $1.45 \times 10^{-5}$<br>$((1.73 \times 10^{-3}) \times (8.38 \times 10^{-3}))$ |
| PP-A +<br>PP-B | $3.38 \times 10^{-4}$ | $6.19 \times 10^{-6}$<br>$((7.39 \times 10^{-4}) \times (8.38 \times 10^{-3}))$ |
| MP-A +<br>PP-A +<br>PP-B | $8.14 \times 10^{-8}$ | $1.07 \times 10^{-8}$<br>$((1.73 \times 10^{-3}) \times (7.39 \times 10^{-4}) \times (8.38 \times 10^{-3}))$ |

#### Simulations of in vitro cocktail PKPD

We simulated the temporal dynamics of bacterial and phage concentrations for single-phage and double-phage cocktail treatments, starting with an initial total phage exposure of 10^4^ PFU/mL (Figure 2A). Consistent with experimental results, single-phage treatments inevitably led to the emergence of resistance by approximately 10 h. In cocktail treatments, MP-A + PP-B and PP-A + PP-B selectively enriched bacteria resistant to MP-A and PP-A, respectively.

**Figure 2.**
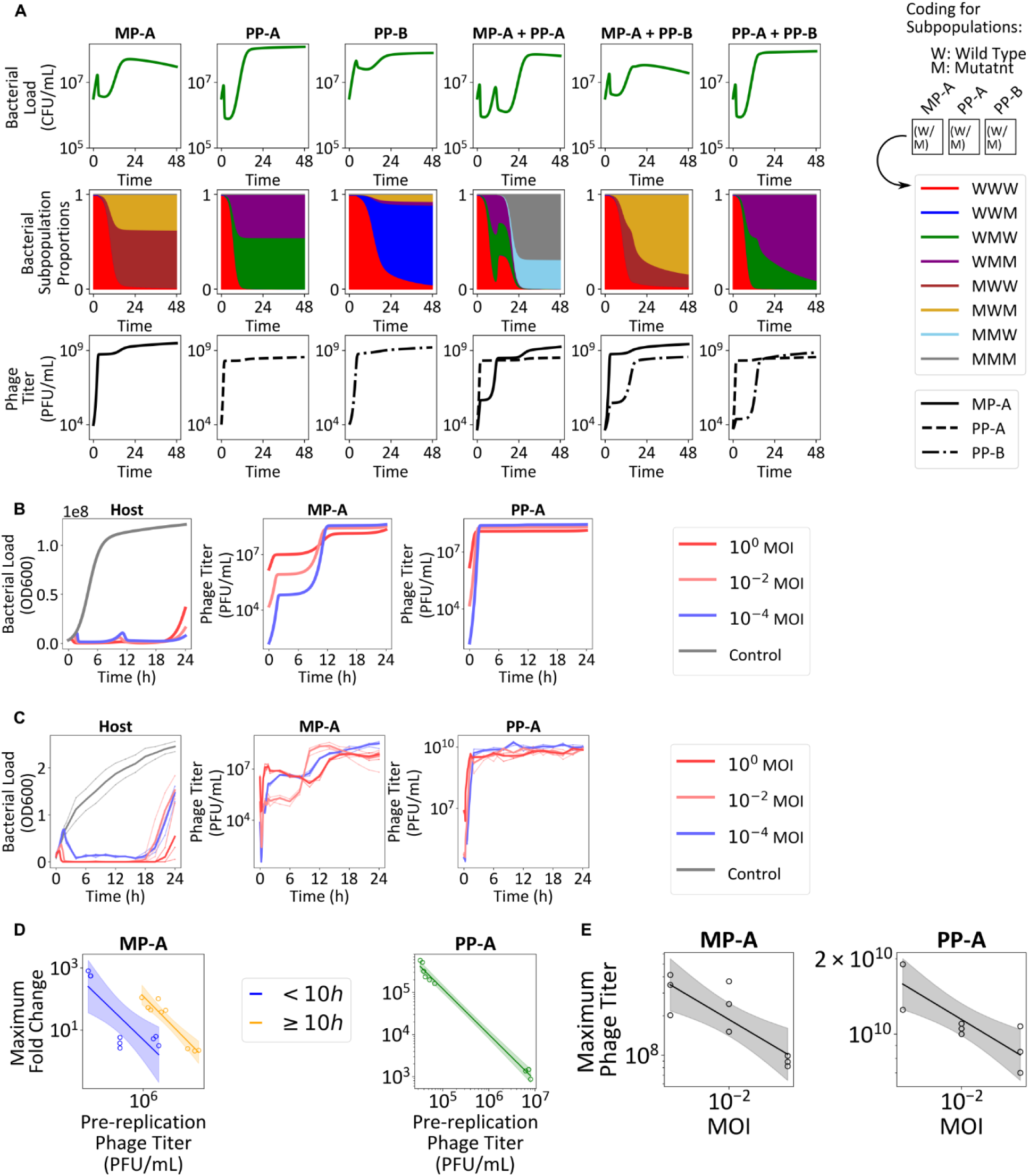
Simulations of cocktail PKPD and *in vitro* validation. (A) The temporal dynamics of bacterial subpopulation fractions were analyzed to evaluate single-phage treatments and double-phage cocktails. Rows 1–3 illustrate the time-course profiles of the total bacterial load, proportions of bacterial subpopulations, and phage titers, respectively. (B) A dose-response simulation of the MP-A + PP-A cocktail at MOI of 0.0001, 0.01, and 1. (C) Bacterial and phage titers were quantified in an *in vitro* PKPD experiment involving the MP-A + PP-A cocktail. The results revealed a dose-dependent suppression of bacterial growth. Initially, PP-A proliferated rapidly, achieving approximately 10^10^ PFU/mL, whereas MP-A showed minimal growth. Between 5 to 10 hours post-treatment initiation, MP-A exhibited a subsequent increase in growth, possibly attributed to predation on emerging PP-A-resistant strains. (D) The magnitude of phage proliferation exhibited an inverse relationship with the pre- replication phage titer. A linear regression line, along with its corresponding 95% confidence interval, is depicted based on log-transformed data. (E) The peak phage titer also demonstrated an inverse correlation with the dosage administered. A linear regression line, along with its 95% confidence interval based on log-transformed data, is presented.

However, substantial fractions of these bacteria were already resistant to PP-B, rendering it ineffective. In contrast, PP-A-resistant bacteria that emerged during MP-A + PP-A treatment included a negligible fraction resistant to both phages, enabling MP-A to effectively suppress these bacteria.

A common pattern observed across all simulations was the initial competition between the two phages, which led to reduced proliferation of the less virulent phage in the early stages. As the population of bacteria resistant to the more virulent phage increased, the less virulent phage exhibited a delayed growth spurt. In MP-A + PP-A treatment, MP-A eventually proliferated to levels similar to those expected under MP-A monotherapy. However, in MP-A + PP-B and PP-A and PP-B treatments, PP-B failed to proliferate to levels expected under PP-B monotherapy owing to the presence of a high number of double-resistant mutants.

A dose-response simulation of the MP-A + PP-A cocktail (Figure 2B) predicted that higher doses, while achieving greater early suppression of bacterial growth through primary infections, inadvertently inhibit secondary infections and reduce phage growth. This prediction is widely supported by the literature^21^ and suggests a potential trade-off associated with higher phage doses.

#### In vitro PKPD experiments with MP-A + PP-A cocktail

We sought to validate the PKPD predictions of the synergistic MP-A + PP-A cocktail *in vitro*. For this purpose, we tracked densities of *P. aeruginosa* 15-4 and concentrations of MP-A and PP-A in the cocktail over 24 hours (Figure 2C). Early treatment phases showed a dose-dependent decrease in bacterial density and a preferential proliferation of PP-A over MP-A. By approximately 8–12 hours of treatment, MP-A demonstrated a secondary growth spurt. By 20 hours, double-resistant bacteria emerged. The replication levels of both phages (Figure 2D) were inversely correlated with their pre-replication concentrations. Notably, replication of MP-A during the first 10 hours was less than that in later stages. Both phage titers peaked inversely to the administered dose (Figure 2E).

Although the MP-A + PP-A cocktail effectively extended the duration of bacterial growth suppression compared to MP-A or PP-A administered alone, multi-phage-resistant bacteria eventually emerged in our experiments.

#### Biofilm elimination assay and model simulation

Biofilm assays were conducted for the two selected phages (MP-A and PP-A) and their combination (MP-A + PP-A) to evaluate their activity against *P. aeruginosa* in a structured biofilm environment (see STAR Methods). A biofilm with an initial OD_600_ of 0.2 was incubated with phages at a total phage concentration of 2x 10^10^ PFU/mL After 6 hours, the mean OD_600_ in the control, MP-A, PP-A, and MP-A + PP-A treated groups was 1.35, 0.49, 0.30, and 0.28, respectively (Figure S6).

To assess whether our model could predict anti-biofilm activity, we first simulated bacterial growth for 24 hours according to the assay protocol and estimated the fraction of dormant bacteria at the end of the simulation. These fractions were then used to set the initial conditions for the next simulation, starting with an OD_600_ of 0.2. However, when applying our original PKPD model to these conditions, the model failed to capture the observed dynamics, likely owing to the structured environment of biofilms and the different fractions of dormant and proliferative bacteria, which can fluctuate depending on various factors.

To address this issue, we modified the model to incorporate spatial heterogeneity, assuming that phage saturation occurs during the infection process.^25^ With this adjustment, the simulations successfully described the experimental results, with predicted OD_600_ values of 1.42, 0.56, 0.17, and 0.16 for the control, MP-A, PP-A, and MP-A + PP-A groups, respectively, at 6 hours (Figure S3B). This modification highlights the adaptability of the model in accounting for different bacterial growth environments and suggests that phage saturation, driven by spatial heterogeneity within biofilms, should be considered when modeling phage therapy in biofilm contexts.

Similar to the time-kill assay results using planktonic bacteria, the model predicted that resistant bacteria would emerge under all conditions. Therefore, we next explored the possibility that, *in vivo*, synergistic interaction of phages with the host immune system could enable successful bacterial control.

### *In vivo* PKPD evaluation

#### Pharmacokinetics of the phage cocktail in noninfected mice

Pharmacokinetic analysis of the MP-A + PP-A phage cocktail in healthy mice across low (10^7^ PFU/mouse), medium (10^9^ PFU/mouse), and high (10^11^ PFU/mouse) doses (Table S2) demonstrated serum half-lives of less than 6 hours for MP-A and 2 hours for PP-A. In contrast, lung half-lives were longer at 10–26 hours for MP-A and 6–19 hours for PP-A. Although the phages were mixed in a 1:1 ratio, PP-A showed initial concentrations 2 to 4 logs higher than MP-A. The concentration-time profiles of both phages are shown in Figures S4A and S4B.

The dose-normalized initial concentrations (c_0_⁄Dase) for PP-A remained consistent in terms of scale, regardless of the dose (Table S2). In contrast, MP-A exhibited a significant increase in terms of scale with escalating doses, indicating a dose-dependent entry of the phage into central circulation from the tail vein. This phenomenon has been previously elucidated using a circulation factor.^26^ The lung-to-serum c ratios were higher in MP-A (0.02–0.17) than in PP-A (0.008–0.01), suggesting better lung penetrance for MP-A than for PP-A.

#### PKPD of the phage cocktail in infected mice

We evaluated the impact of escalating intranasal doses of *P. aeruginosa* on host survival. Mortality rates increased significantly beyond a bacterial dose of 10^7^ colony-forming units (CFU)/mouse, with a critical decline in survival at 5x 10^7^ CFU/mouse (Figure S5), indicating a tipping point for the host immune response.^27,28^

To ensure both adequate PKPD sampling and successful induction of pneumonia, we selected a bacterial dose of 10^7^ CFU/mouse for subsequent experiments. Phages were administered as intravenous (i.v.) bolus injections one hour after infection, and concentrations of phage and bacteria were measured at pre-specified time points from both the serum and lungs.

The infected-to-noninfected lung Auc_last_ ratios of MP-A were 6.98 × 10^5^, 517.16, and 2.40 in the low, medium, and high dose groups, respectively, whereas corresponding values for PP-A were 6.96 × 10^5^, 1.39 × 10^4^, and 73.33, respectively (Table S3). This observation indicates substantial phage replication of both MP-A and PP-A in the presence of target bacteria and suggests that phage replication inversely correlates with the administered dose and is greater for PP-A than for MP-A.

Phage treatment led to a greater early reduction of bacterial load compared to the control, which increased in a dose-dependent manner. However, a gradual resurgence in bacterial load was observed after 12 hours across all phage doses. In the medium and high dose groups, bacterial concentrations eventually approached approximately 10^4^ CFU/mL. The PKPD profiles for both serum and lung concentrations are shown in Figures S4A, S4B, and S4C.

Survival curves (Figure S4C) showed significant improvements in the medium (10^9^ PFU/mouse) and high (10^11^ PFU/mouse) dose groups within 60 hours relative to the control (p=0.006), whereas the low dose (10^7^ PFU/mouse) group did not show a significant difference from the control (p=0.3).

### Extended mathematical model incorporating phage PK and immune system dynamics

We extended our mathematical model to incorporate both *in vivo* phage PK and host immune response dynamics. A two-compartment model with linear elimination was employed to capture the exposure-time profiles of phages within the lung environment. We assumed that immune cells were activated by bacteria in the P state, enabling them to kill bacteria in this state but not those in the D state. Building on the work of Leung and Weitz et al., we posited that both immune activation and immune-mediated bacterial clearance become saturated at high bacterial concentrations.^29^ This refined model, illustrated in Figure 3A, accounts for the complex interactions between phage activity and the host immune response. We calibrated the model parameters within a physiologically relevant range based on observed *in vivo* PKPD measurements. Compared to the rate *in vitro*, the *in vivo* rate of dormant state transition was approximately 10 times lower, resulting in a higher maximum bacterial density. In addition, the *in vivo* per bacterium infection rate of PP-A was approximately 70% reduced, whereas its apparent burst size was 8 to 9 times increased. The predictions of the model showed a reasonably good fit with the experimental observations (Figure 3B).

**Figure 3.**
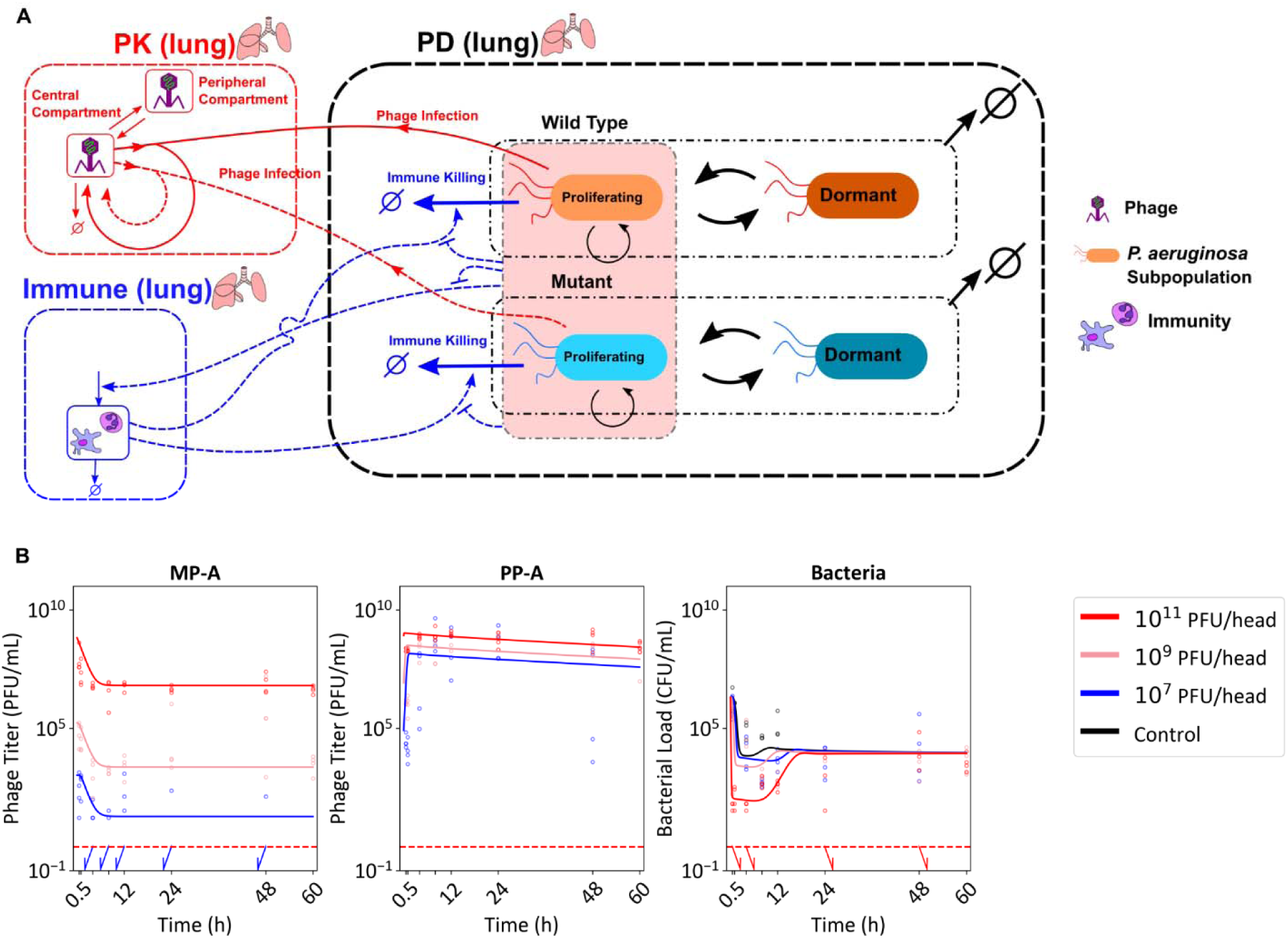
Extension and calibration of the *in vitro* PKPD model to predict *in vivo* PKPD. (A) The schematic representation of the extended *in vivo* PKPD model incorporates both phage PK and the immune response. (B) The predictions generated by the calibrated in vivo PKPD model are superimposed on the empirical observations.

#### Dose response of MP-A + PP-A and PP-A under varying immune status

We first assessed the effects of different initial bacterial loads in the absence of phage treatment. As observed earlier, a critical threshold of approximately 10^6.8^ CFU/mL marked the sharp transition from effective immune-mediated bacterial clearance to uncontrolled bacterial growth.

With phage treatment, bacterial load consistently decreased and converged towards 10^4^ CFU/mL, irrespective of the initial bacterial load. In addition, phage treatment reduced the extent of immune activation (Figure 4A).

**Figure 4.**
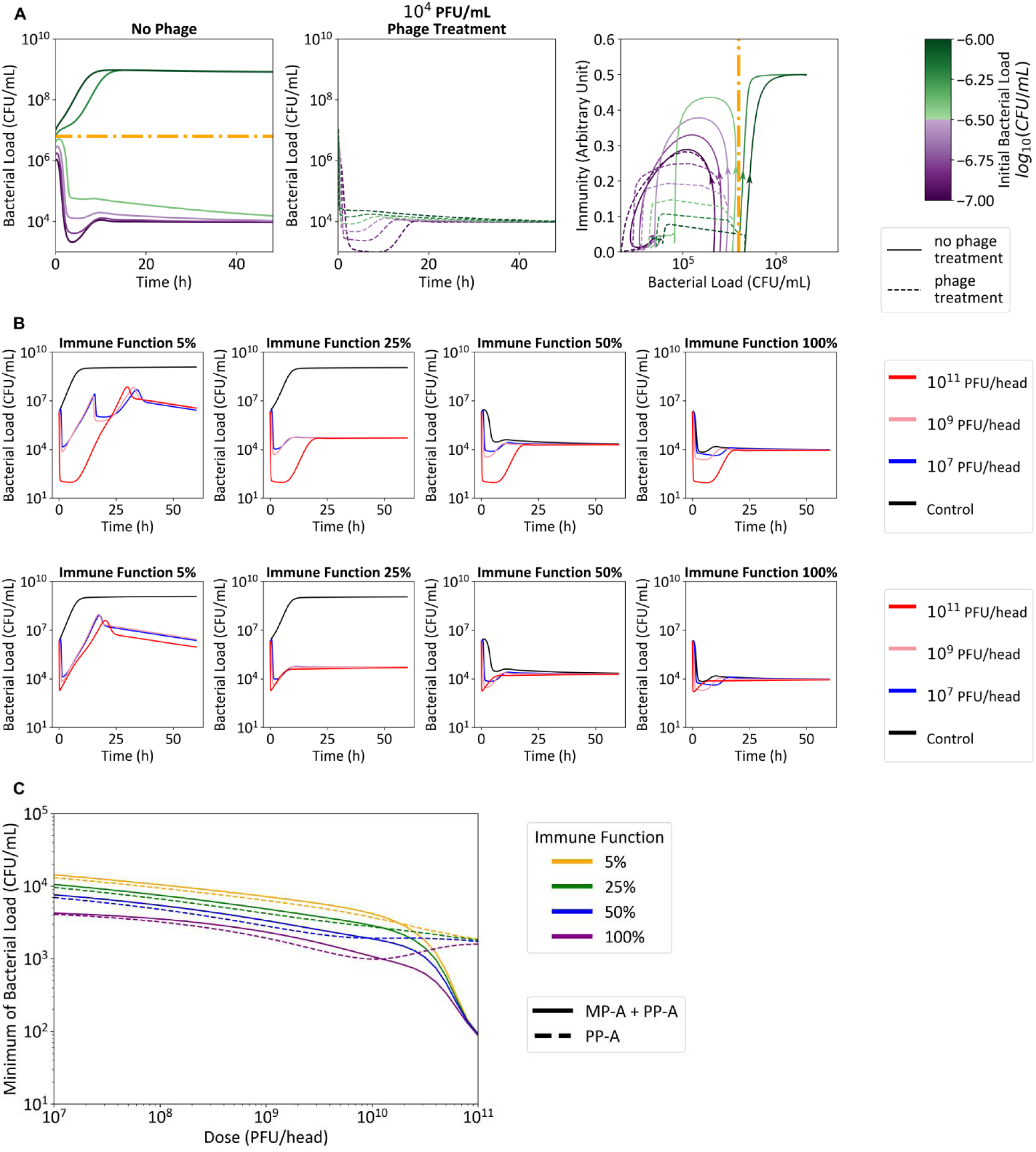
Phage synergy with immunity. (A) Simulations to analyze the clearance of bacterial load in relation to the initial bacterial load, both in the presence and absence of bacteriophages. (B) The temporal dynamics of bacterial clearance that were examined for PP-A monotherapy and the combined MP-A + PP-A therapy, with outcomes contingent upon varying levels of immune function. (C) The minimum bacterial load was evaluated as a function of phage dosage and immune function for PP-A monotherapy and MP-A + PP-A combination therapy.

We next explored the impact of phage doses for the MP-A + PP-A cocktail and PP-A monotherapy under varying immune statuses (Figure 4B). Results indicated that the bacterial load at 60 hours was primarily determined by host immunity and tended to decrease with immune function for both treatments. Higher phage doses were associated with a greater early reduction in bacterial load. The cocktail therapy performed better than monotherapy in achieving early bacterial reduction at doses above 10^10^ PFU/head, but both treatments were comparable at doses below 10^10^ PFU/head (Figure 4C).

These results suggest that higher phage doses enhance the speed of bacterial load reduction while inhibiting excessive immune activation. However, phage cocktails provided no substantial benefit over single-phage treatments unless the administered dose exceeded a critical threshold. A minimum level of host immunity is necessary for successful phage therapy.

### *In vivo* model validation

#### Comparative in vivo efficacy

Having developed the full version of our model that incorporated *in vivo* factors, we finally sought to validate its predictions. We induced acute pneumonia in mice by intranasally inoculating a higher bacterial dose of 10^7.7^ CFU/mouse of *P. aeruginosa* 15-4. Treatment groups received individual phages (MP-A, PP-A) and a cocktail (MP-A + PP-A mixed in equal ratios) at total doses of 5 x 10^9^ PFU/head as i.v. boli. At 21 hours post-infection, all animals were sacrificed to measure lung bacterial loads. All treatments significantly reduced bacterial loads compared to controls (10^8.52^ CFU/lung), with titers of 10^5.18^, 10^4.77^, and 10^4.76^ CFU/lung for MP- A, PP-A, and MP-A + PP-A, respectively. While the reduction in bacterial load of the treatment groups compared to the control was significant, no significant difference was observed between the treatment groups (Figure 5A). This result was contrary to the *in vitro* results in which the phage strains exhibited different efficacies and the cocktail was more effective in preventing regrowth of resistant bacteria (Figure S2A).

**Figure 5.**
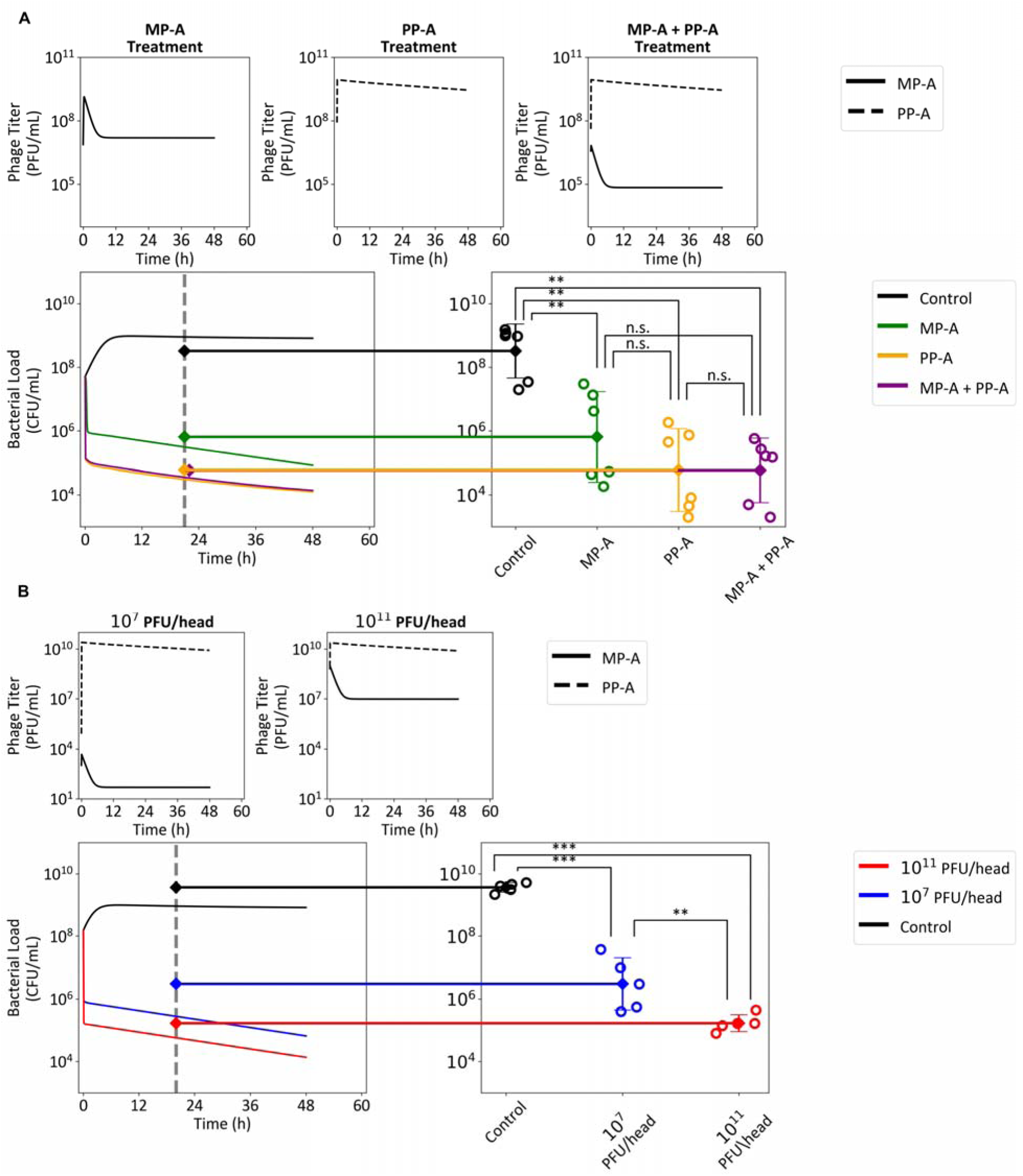
Validation of the *in vivo* PKPD model. The model validation involved assessing the comparative *in vivo* efficacy of monotherapy and cocktail therapy (A) and dose-response relationships (B) in mice with acute pneumonia. The top panels of (A) and (B) display the predicted phage PK. The bottom left panels of (A) and (B) show the predicted bacterial density over time, whereas the bottom right panels present the observed bacterial densities measured at 21 (A) and 20 (B) hours post-infection. The error bars represent the mean ± standard deviation for the log10-transformed data. Analysis of variance (ANOVA) indicated significant differences, which were further examined using Tukey’s honestly significant difference (HSD) test. Statistical significance is denoted as follows: not significant (n.s.), **p<0.01, ***p<0.001.

Model simulations, assuming the same bacterial dose of 10^7.7^ CFU/mouse and a typical lung volume of 1 mL in mice^30^, reasonably mimicked the experimental findings. Specifically, our model successfully reproduced the lack of substantial benefit of the MP-A + PP-A cocktail over PP-A monotherapy at 5 x 10^9^ PFU/head dose (Figure 5A).

#### In vivo dose-response relationship

We conducted a dose-response study of MP-A + PP-A using a higher bacterial inoculum of 10^8.17^ CFU/mouse to validate the extrapolative capability of the model. At 20 hours, bacterial loads were 10^9.56^, 10^6.52^, and 10^5.19^ CFU/lung in the control, low dose (10^7^ PFU/head), and high dose (10^11^ PFU/head) groups, respectively. While the model predictions did not match the observations exactly, they reasonably described the overall dose-dependent reduction in bacterial load (Figure 5B).

A similar, although less pronounced, dose dependence was observed in a subsequent survival experiment involving an acute pneumonia mouse model inoculated with 5 x 10^8^ CFU/head of *P. aeruginosa* 15-4. In the control group, 3 out of 8 mice survived by day 4 (Figure S6). In the groups receiving 5x 10^4^, 5x 10^6^, 5x 10^8^, and 5x 10^10^, PFU/head, 5, 5, 7, and 6 out of 8 mice survived, respectively (Figure S6). This result showed that host survival tended to improve with higher phage doses.

### Stability Testing of Selected Phages

Stability assays were conducted for the two phages selected for further development (MP-A and PP-A) to assess their long-term viability under different storage conditions (see STAR Methods for details). PP-B was excluded from further development owing to its limited efficacy and compatibility.

#### Thermal stability

The thermal stability of MP-A and PP-A was evaluated across a range of temperatures (4°C, 25°C, 37°C, 50°C, 60°C, and 70°C) over a 60-minute period (Figure S7A,B). MP-A retained its viability at temperatures up to 50°C, with approximately 84% of the initial titer remaining after 60 minutes at this temperature. However, at 60°C and 70°C, the titer declined more substantially, with only 79% and 53% remaining, respectively. PP-A showed greater susceptibility to higher temperatures, retaining only 24% of its titer at 60°C and 1% at 70°C after 60 minutes of exposure (Figure S7A,B). These results suggest that MP-A is more thermally stable than PP-A, indicating that MP-A may be more suitable for applications requiring exposure to elevated temperatures.

#### pH Stability

The pH stability of MP-A and PP-A was also tested across a broad pH range (pH 2–10) over a 30-day period. MP-A demonstrated remarkable stability, with titers remaining relatively consistent (≥70% of the initial titer) across a neutral to mildly alkaline pH range (pH 6–9) (Figure S7C,D). However, phage viability rapidly decreased under acidic conditions, with no detectable phage activity by day 7 at pH 2. PP-A displayed a similar pH stability profile, with significant reductions in titer occurring at pH 2 and pH 3. Both phages were most stable in the neutral pH range, with greater than 90% of their original titer remaining after one day of incubation at pH 7.

## Discussion

In this study, we conducted a comprehensive evaluation of the PKPD for three phages that target *P. aeruginosa* and assessed their combinatorial effects. Our integrated approach combined *in vitro* experiments, *in vivo* models, and *in silico* simulations. Our *in vitro* studies revealed that phage resistance inevitably develops during mono-phage treatments, which typically emerges around 10 hours post-phage inoculation. Among the tested phage combinations, only the MP-A + PP-A cocktail successfully mitigated resistance. A model-based analysis revealed that this cocktail was successful owing to the collateral sensitivity between MP-A and PP-A. However, even this effective cocktail eventually succumbed to multi-phage-resistant bacteria. Compared to *in vitro* observations, our *in vivo* studies revealed a different trend. The PKPD experiments in mice showed that phage treatments reduced bacterial loads in a dose-dependent manner, and the growth of phage-resistant bacteria was minimal. At phage doses of 10^9^ and 10^11^ PFU/mouse, bacterial load converged towards approximately 10^4^ CFU/mL. Model simulations indicated that an intact immune system *in vivo* can effectively control the growth of phage-resistant bacteria. Higher phage doses lead to a greater early reduction of bacterial load and reduce the extent of immune activation. However, when the host immune function falls below a critical threshold, phage-resistant bacteria can no longer be suppressed and thus grow unchecked. In such conditions, alternative strategies, such as inclusion of additional phages, may be needed to achieve complete coverage of all bacterial mutants.

Phage therapy exhibits unique PKPD properties owing to the self-replicating nature of phages. Unlike conventional antibiotics, the effectiveness of phage exposure depends not only on the administered dose but also on the replicative capacity of the phage and the density of target bacteria. This complexity has led to the application of mathematical modeling to predict phage therapy outcomes.^31–35^ Cairns et al. reported that a simple mathematical model provided a surprisingly good fit to *in vitro* experimental data.^19^ However, Weld et al. showed that similar models poorly predicted *in vivo* outcomes.^36^ Roach et al. found that bacterial clearance predicted by an earlier mathematical model overestimated *in vivo* observations.^25^ They addressed this issue by incorporating spatial heterogeneity and phage saturation effects in their model. Recently, Delattre et al.^20^ directly fitted their model to *in vivo* data, using *in vitro* data primarily to estimate the median eclipse phase and the burst size of the phage. In this study, we noticed significant differences between *in vitro* and *in vivo* conditions, particularly in terms of maximum bacterial density and PP-A parameters. These findings align with current evidence suggesting that mathematical models based solely on *in vitro* data have clear limitations for *in vivo* extrapolations. Integrating *in vitro* and *in vivo* data during model development can bridge this gap and provide a more comprehensive understanding of phage therapy efficacy.

Our work proposes important principles for cocktail and dose optimization. In the absence of an immune response, it is important to combine phages with low cross-resistance to ensure adequate therapeutic depth.^37^ One of the possible causes of poor phage therapy outcomes in immunocompromised individuals^22^ is the inability to control the emergence of phage-resistant bacteria. Furthermore, higher doses may inadvertently promote resistance owing to reduced phage replication. For example, Chang et al. reported that in neutropenic mice, higher phage doses led to decreased phage proliferation and increased phage-resistant bacterial populations.^21^ In immunocompetent hosts, phage resistance emergence is less problematic.^25,38,39^ The host immune system can effectively scavenge minor phage-resistant mutants, reducing the need for high cocktail depth. This is also supported by our experimental result, where the MP-A + PP-A cocktail demonstrated negligible enhancement in bacterial load reduction compared to PP-A monotherapy at a dose of 5 x 10^9^ PFU/head (Figure 5A). We attribute this to the immune system compensating for the role of MP-A by effectively eliminating PP-A resistant mutants. However, our simulations (Fig 4B) predicted that at higher doses, the MP-A + PP-A cocktail would achieve greater early reduction. This effect is due to MP-A’s contribution in targeting minor populations of PP-A resistant mutants during primary infection.

Since higher doses are generally favorable in immunocompetent hosts, optimizing the administration route to maximize the initial phage exposure at the site of infection is also crucial. For example, direct inhalation of phages in pneumonia cases may yield higher early exposure than alternative routes of administration (e.g., intravenous or oral). From a practical perspective, *P. aeruginosa* is a major cause of nosocomial infections, particularly in critically ill or immunocompromised patients.^40^ According to our simulations (Figure 4B), an immune function of below 25% in healthy hosts was associated with an outgrowth of phage-resistant bacteria. Similarly, a cutoff of 20% was proposed for the minimum level of neutrophil activity.^25^ A clear treatment guideline for immunocompromised patients should be established through clinical studies in future. Enhancing immune function through adjunctive therapies could improve phage therapy outcomes.

Our findings also highlight that phage life history traits such as adsorption rate, latent period, and burst size measured in isolation through adsorption assay and one-step growth assay, respectively, have limitations in predicting phage efficacy. For instance, adsorption assay suggested that PP-A exhibits incomplete adsorption, with a significant fraction of unadsorbed phages remaining after 30 minutes, and one-step growth assay suggested that it has the lowest burst size (∼20 per infection cycle). However, time-kill assay showed that it demonstrated the strongest early virulence, which may be due to its short latent period. Apparent adsorption rate constants estimated using our model corroborated these observations, showing the highest value for PP-A, followed by MP-A and PP-B. Due to the simplifying model assumption of immediate lysis of infected bacteria that disregards variations in latent periods, the estimated infection rate constants represent composite effects of both adsorption rates and latent periods. Additional simulations using a model that explicitly incorporated latent periods (data not shown) confirmed that PP-A’s significantly shorter latent period was primarily responsible for its superior efficacy. Hence, to effectively link phage life history traits to bacterial lytic efficiency, these traits must be integrated into a mathematical model. Alternatively, their composite effects could be estimated using a simplified model as that proposed in our study.

Another important challenge is to explore phage efficacy in patients with chronic *P. aeruginosa* infection, in which bacteria exist primarily in a biofilm state. Our experimental results indicated reduced efficacy in biofilm conditions compared to that in planktonic conditions. While our original model accurately predicted planktonic bacterial dynamics, it required modifications to adequately reflect the biofilm environment. By incorporating phage saturation into the infection process, we could account for the structured nature of biofilms, aligning the model predictions with the experimental observations. Our findings demonstrate the importance of considering spatial heterogeneity when modeling phage therapy in biofilms and suggest that this might be achieved through relatively simple adjustments of the model.

The flexibility of our model was further demonstrated through exploratory simulations that addressed two clinically relevant scenarios: the efficacy of phage treatment against stationary-phase bacteria and the potential synergy between phages and antibiotics (Figure S8A-D). When phages were administered after bacteria had reached the stationary phase, bacterial reduction was significantly slower (Figure S8A). This suggests that phage therapy may be less effective in established infections wherein bacterial growth has stabilized.

To explore combination therapies, we simulated the co-administration of a hypothetical antibiotic with the phages. The simulations revealed that if an antibiotic that can target both dormant and proliferating bacteria is used in combination with the phage(s), the problem of low efficacy of phage therapy against stationary-phase bacteria can be overcome (Figure S8A). This synergy was particularly evident when antibiotics were administered in multiple doses, demonstrating the potential for phage-antibiotic combinations to target stationary-phase bacteria more effectively.

Varying the ratios of individual phages within a cocktail could potentially influence bacterial killing efficiency. Stratifying the time-kill assay results by mixing ratio (Figure S9) suggested that differences among various mixing ratios were minimal. However, mixing ratio may be important when using a more unbalanced mixing ratio (such as 1:100) than that used in our study or for phage strains not used in this study. There is need for future research to determine the optimal mixing ratios of phage cocktails, particularly under different dosing regimens and treatment scenarios.

Despite the advancements presented in our study, certain limitations of the work must be acknowledged. Our research heavily relied on mathematical modeling and experiments using specific phage strains against *P. aeruginosa*. Phage-bacterial interaction dynamics can vary significantly across different systems, which may affect the generalizability of our results.

Secondly, although the model incorporated immune response dynamics to explain the differences between *in vitro* and *in vivo* findings, we did not explicitly assess immune biomarkers. Thus, the model may not fully represent the complexities of immune interactions. The change in the actual immune cell profiles upon phage therapy and at variable doses will be relevant to examine directly in future studies. For example, a recent report^41^ revealed that alveolar macrophages can directly reduce phage titers and that depleting these macrophages prior to phage treatment resulted in reduced pulmonary bacterial loads. This study suggests that, beyond the competition between phages and immune cells to reduce bacteria, the immune-mediated clearance of phages must also be considered—an aspect our current framework does not yet include. Additionally, our study did not assess *in vivo* efficacy in immunocompromised hosts, making our model predictions for these hosts speculative. Lastly, while our model has provided valuable insights, further validation through clinical trials is necessary to translate the findings to clinical settings. The effectiveness of phage therapy in humans, particularly among diverse patient populations, needs to be rigorously tested.

In conclusion, our study represents a significant advance in understanding phage PKPD. By integrating *in vitro*, *in vivo*, and *in silico* methodologies, we have not only evaluated phage therapy comprehensively but also developed a predictive model capable of generating testable hypotheses. Most crucially, our model has illuminated key principles for optimizing phage cocktails and doses based on host immune status. The insights gained from our research lay the foundation for future studies and pave the way for more effective phage-based treatments.

## Acknowledgements

This research was supported by a grant (No. HI23C121600) from the Korea Health Industry Development Institute (KHIDI) acquired by D.Y. and D.C. This research was additionally supported by a grant (No. P0023226) from KHIDI and the Brain Korea 21 FOUR Project for Medical Science, Yonsei University College of Medicine, Seoul, Republic of Korea.

## Author contributions

D.Y. and D.C. conceived the project. K.K., H.J.Y., D.K., H.H.P., and D.Y. designed and conducted the experiments. J.S.C. and D.C. performed PKPD modeling and data analysis. J.S.C. and D.C. wrote the original draft. C.O.K. and B.H.J. reviewed and edited the draft.

## Declaration of generative AI and AI-assisted technologies in the writing process

During the preparation of this work, the authors utilized ChatGPT in order to check for grammatical errors and make the tone of the manuscript more formal and academic. All products of the tool were reviewed and re-edited by the authors. The authors bear complete responsibility for the content of the published article.

## STAR Methods

### RESOURCE AVAILABILITY

#### Lead contact

Further information and requests for resources and reagents should be directed to and will be fulfilled by the lead contact, Dongwoo Chae.

#### Materials availability

Bacterial strains and phage strains are available only with permission from Microbiotix.

#### Data and code availability

- All data analyzed in this paper were deposited to Mendeley Data and is publicly available as of the date of publication.
- All code used in this study has been uploaded to Mendeley Data and is publicly available as of the date of publication.
- Any additional information required to reanalyze the data reported in this paper is available from the lead contact upon request, if permission is granted by Microbiotix.

### EXPERIMENTAL MODEL DETAILS

#### Bacterial strains and phages

##### Phage isolation and preparation

Phages MP-A, PP-A, and PP-B were isolated from sewage samples collected at Severance Hospital, Seoul, Korea. Samples were treated with 1 M NaCl and refrigerated at 4°C for 24 hours. Following centrifugation at 8000 *g* for 30 minutes at 4°C (Supra R22; Hanil, Korea), the supernatant was filtered through a 0.22-μm pore-size membrane (Corning, NY, USA).

Polyethylene glycol (PEG) 8000 at 10% concentration (Sigma, St. Louis, MO, USA) was added and the mixture was incubated for another 24 hours at 4°C. Phages were pelleted by centrifugation at 14000 *g* for 1 hour at 4°C, resuspended in sterilized SM buffer (100 mM NaCl, 8 mM MgSO_4_, 50 mM Tris-HCl, pH 7.5), and further sterilized by passage through a 0.22 μm syringe filter.

##### Host strain and maintenance

The *P. aeruginosa* 15-4 strain was isolated from clinical samples of a pneumonia patient from Severance Hospital. Its resistance to carbapenems (imipenem and meropenem) was confirmed using the VitekN132 system (bioMérieux, Marcy-l’Étoile, France) and CLSI disk diffusion method. The strain was maintained as a frozen stock in Luria broth LB medium with 25% glycerol at - 70°C and reactivated for experiments by overnight incubation at 35°C in LB broth on a rotary shaker at 180 rpm.

##### Phage-sensitivity testing

Sensitivity of *P. aeruginosa* 15-4 to the phages was assessed using the spot test on Mueller-Hinton II agar (MHA) plates (Asanpharm, Seoul, Republic of Korea). The host bacteria were spread evenly on the agar plates; phages were spotted, followed by serial dilution and incubation using the double-layer agar method. Single-plaque phages were harvested, amplified in LB medium with the host strain, and cleared of cell debris by centrifugation and filtration. The phage preparations were then stored in 1.5-mL aliquots at 4°C.

#### Mouse acute pneumonia model

The experimental protocol for the mouse acute pneumonia model was approved by the Institutional Animal Care and Use Committee of the respective institutions where the experiments were conducted (HDS Bio, Pohang-si, Republic of Korea, IACUC ID: 20231024-21; DT&CRO, Yongin-si, Republic of Korea, IACUC ID: 23E019; and HLB Biostep, Incheon, Republic of Korea, IACUC ID: 24-HB-0126).

For the *in vivo* validation experiments, five-week-old male ICR mice were used to compare lung bacterial loads following MP-A and PP-A monotherapy with those after administration of the MP- A + PP-A cocktail, all administered at the same dose. To compare different doses of the MP-A + PP-A cocktail, six-week-old female mice were used. The mice were housed in polycarbonate cages, under a temperature of 19℃ and relative humidity of 50–60%, with free access to food (Teklad Certified Irradiated Global 18% Protein Rodent Diet 2918C, Envigo RMS, Inc., IN, USA).

In the *in vivo* survival experiments evaluating dose-dependent responses to phage treatment, seven-week-old female ICR mice were employed. Additionally, six-week-old male ICR mice were used for both the dose-ranging survival experiments and the *in vivo* PKPD studies. The mice were housed in polycarbonate cages, under a temperature of 20–26℃ and relative humidity of 40–70%, with free access to food (Rodent Diet, Purina®, Neenah, WI, USA).

CO_2_ was used to euthanize all animals.

##### Preparation of *P. aeruginosa* for inoculation

The *P. aeruginosa* 15-4 strain was prepared using the following protocol. The bacteria stored at -80°C were smeared on MHA and incubated at 37°C for 24 hours. Two colonies were picked from the MHA, one for establishing the ratio between optical density (OD) and CFU and the other for inoculation into the mice. The colonies were each inoculated in the Mueller-Hinton II Broth (MHIIB). The inoculated broth was incubated in a shaking incubator at 150 rpm for 24 hours at 37°C. The media was centrifuged at 15,000 rpm (21206 x g), for 2 minutes. The pellet was resuspended in 1 ml of autoclaved normal saline solution. For the first colony, the resuspended bacterial solution was diluted by 10-fold and the OD was measured using a spectrophotometer (Epoch2, Biotek, Seoul, Korea). Subsequently, 100 μL of the mixture was smeared on MHA and incubated at 37°C overnight. The number of colonies was compared to the OD_600_ to calculate the concentration of bacteria in the mixture. For the second colony, based on the OD_600_ measured by the spectrophotometer, the mixture was diluted to the target levels to be used for inoculation into mice.

##### Inoculation of *P. aeruginosa* 15-4 into mice

A total of 20 μL of the *P. aeruginosa* 15-4 mixture prepared above was inoculated intranasally into the mice 2 hours before injection of the phage treatments into the tail vein.

### METHOD DETAILS

#### Bacteriophage characterization

##### Genomics analysis

Genomic DNA was extracted from the phage stock using the Phage DNA Isolation Kit (Norgen Biotek, ON, Canada) and quantified using the QuantStudio 3 (Thermo Fisher Scientific, MA, USA) system. The prepared DNA was then used for library construction for whole-genome sequencing (WGS) using the TruSeq Nano DNA Kit (Illumina, CA, USA). Sequencing was performed on an Illumina sequencing platform based on Sequencing by Synthesis (SBS) technology.

Raw sequencing reads were processed to remove adapters and low-quality bases using Trimmomatic v0.36.^42^ Quality-filtered reads were assembled de novo using SPAdes v3.15.0.^43^ The resulting contigs were annotated using Prokka v1.14.6^44^ and compared to reference databases (NCBI BLAST) to identify homologous sequences. Genome coverage and completeness were assessed using QUAST v5.0.2.^45^ Phage genomes were visualized and analyzed for key genes related to phage structure, replication, and host interaction. Annotation of tRNAs was performed using tRNAscan-SE v2.0.7.^46^

##### Electron microscopy

Phage morphology was visualized using transmission electron microscopy (TEM). Purified phage suspensions were deposited onto Formvar/carbon-coated 200 mesh copper grids (TED PELLA, INC., CA, USA) and allowed to adsorb for 15 seconds. Excess liquid was removed with 110 mm filter paper (ADVENTEC, Tokyo, Japan), and the grids were negatively stained with 1% (w/v) uranyl acetate for 30 seconds. The grids were air-dried and examined using HT7800 TEM (HITACHI, Ltd., Tokyo, Japan) at the Yonsei Biomedical Research Institute, Yonsei University college of medicine.

Micrographs were captured using OneView 4K digital camera (Gatan, Inc., CA, USA). Morphological features, including head diameter, tail length, and tail type (contractile or non-contractile), were measured from the images using ImageJ software v1.53t.^47^ Phages were classified according to the International Committee on Taxonomy of Viruses (ICTV) guidelines into the appropriate families (e.g., Myoviridae, Podoviridae).

##### Phage adsorption assay

A single colony of host bacteria (MXB1001, PAE 15-4) was cultured in 5 mL of LB medium at 35°C with shaking (180 rpm) for 16 hours. This pre-culture was then expanded into 20 mL of LB medium until reaching an OD_600_ of 0.5. The bacterial cells were mixed with phages MP-A, PP-A, or PP-B with MOI of 10^-3^, vortexed, and filtered through a 0.22-μm pore-size syringe filter.

Samples (1 mL) were collected at intervals from 0 to 30 minutes and analyzed for phage concentration using the double agar overlay method. The experiments were performed in triplicate.

##### One-step growth assay

Similar to the adsorption assay, a single colony of MXB1001 (PAE 15-4) was incubated in 5 mL of LB medium at 35°C with shaking (180 rpm) for 16 hours. Subsequently, 500 μL of this culture was transferred into three 5 mL tubes of LB medium and incubated until an OD_600_ of 0.2 was reached.

Bacteria were mixed with phages with MOI of 10^-3^ for MP-A and 10^-5^ for PP-A and PP-B. The mixture was incubated for 10 minutes at 18–29°C and then centrifuged at 10,000 rpm (9,400 x g) for 10 minutes; the supernatant was discarded, and pellets were resuspended in 1 mL of LB medium. This washing step was repeated once. Twenty microliters from the final suspension were inoculated into 20 mL of LB medium and incubated at 37°C with shaking (180 rpm).

Samples (100 μL) were taken every 5 minutes for 2 hours and processed using the double agar overlay method to quantify phages. The experiments were performed in triplicate.

#### In vitro bacterial time-kill assay

##### Conventional high-throughput assay

1. Preparation: A single colony of host bacteria (MXB1001, PAE 15-4) was cultured in 5 mL of MHIIB in a 15-mL conical tube and incubated at 35°C with shaking (180 rpm) for 16 hours.
2. Inoculation: 100 μL of this culture was transferred into 9.9 mL of MHIIB in a 50-mL conical tube and incubated until it reached an OD_600_ of 0.12 (equivalent to approximately 1.00IZJ10^8^ CFU/mL).
3. Assay Setup: The assay was set up in a 96-well plate with a total volume of 100 μL per well. Bacteria were added to each well to achieve a starting concentration of 5.0 x 10^6^ CFU/mL). Phage treatments (both single and cocktail) were added at 50 μL per well to achieve designated MOI values (ranging from 10^-6^ to 10^2^); replicate numbers were recorded and are provided in Table S4.
4. Incubation and Measurement: Plates were incubated at 35°C for 24 hours in a Spectramax ABS spectrophotometer (Molecular Devices, San Jose CA, USA), with OD_600_ measurements taken hourly after shaking the plate.

##### Extended assay

This extended assay simultaneously measured bacterial load and phage titer.

1. Preparation: A single colony was cultured in 10 mL of LB medium in a 15-mL conical tube at 35°C with shaking (180 rpm) until an OD_600_ of 0.2 was reached.
2. Dilution and Inoculation: 2 mL of this culture was diluted into 20 mL of LB medium. Phages were added to achieve the specified MOI. The control group consisted of two replicates. There were three treatment groups for which the MOI was 10^-4^, 10^-2^, and 10^0^. Six replicates were conducted for each treatment group.

Incubation and Sampling: Single-phage treatments were incubated for six hours, with OD_600_ measurements taken hourly. The MP-A + PP-A cocktail was incubated for 24 hours, with hourly measurements of both OD_600_ and phage titers. Phage titers for MP-A were measured using PAE 14-4 (resistant to PP-A) and for PP-A using PAE 51540 (resistant to MP-A), employing the double agar overlay method. Conversion factors of 1/0.9 for MP-A and 1/0.4 for PP-A were applied to adjust plaque counts to levels equivalent to those obtained with PAE 15-4.

#### Phage therapy population dynamics model

In the development of the population dynamics model, we employed a deterministic approach using ordinary differential equations (ODEs). We followed the standard mass action kinetics framework, assuming that phages and bacteria are evenly distributed within a well-mixed environment and their concentrations can be approximated using a continuous scale.

The following equations describe the population dynamics of phage-bacteria interactions under mono-phage treatment that yielded the best fit to our data. The definitions of all the model parameters are given in Table 1.

(Wild-type bacteria)

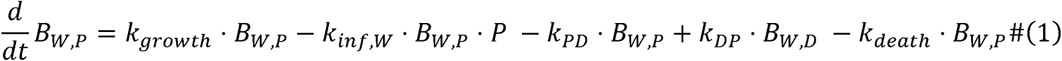

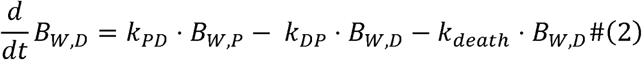

(Resistant mutants)

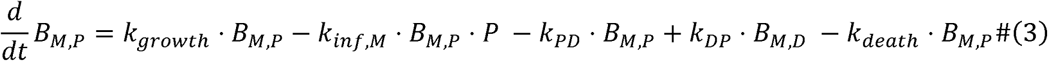

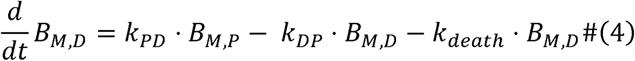

(Density dependence of dormant state transition)

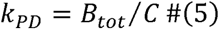

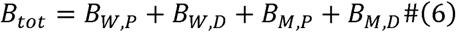

(Phages)

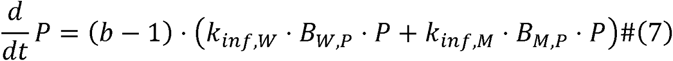

(*B_W,P_* Wild-type, proliferating bacteria, *B_W,D_*: Wild-type, dormant bacteria, *B_M,P_*: Mutant, proliferative bacteria, *B_M,D_*: Mutant, dormant bacteria, *P*: Phages)

The initial condition of the system was as follows:

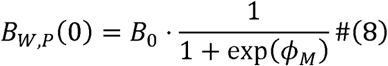

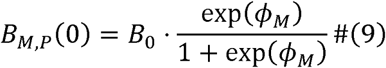

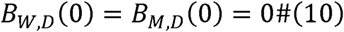

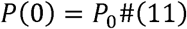

To extend the model to cocktail therapy, all possible phage-sensitivity combinations with regards to M,.P-,A, PP-A, and PP-B were considered. A three-digit subscript notation, *B_ijk_*, with i,j,k ∈ {*W*,*M*}, was used to represent each bacterial subpopulation. The proliferating or dormant state was additionally distinguished using an additionally indicator symbol of either P or D following a comma separator. Thus, the total bacterial concentration is a sum of all possible genetic variants in both P and D states:

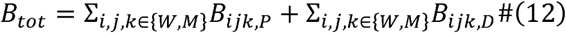

To reduce unnecessary clutter in the parameter symbols, we assigned integer identifiers of 1, 2, and 3 to phages MP-A, PP-A, and PP-B, respectively. Accordingly, the infection rate constants for MP-A, PP-A, and PP-B of the *B_i,j,k,P_* (*i,j,k* ∈ {W,M}), and *k_iinf,1_* (*i,j,k)*, *k_iinf,2_* (*i,j,k)* and *k_iinf,3_* (*i,j,k)* respectively. Hence, for θ ∈ {1,2,3},

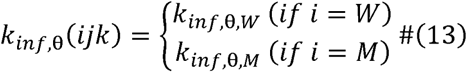

The following equations describe the dynamics of different bacterial variants, where *i,j,k* ∈ {*W*,*M*}

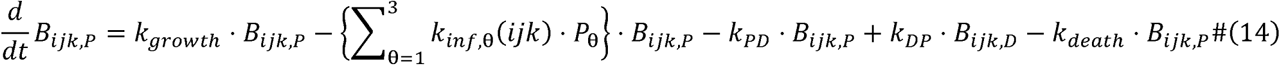

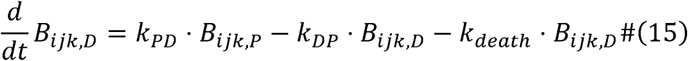

The dynamics of phages are described by the following equations, where *θ* ∈ {1,2,3}

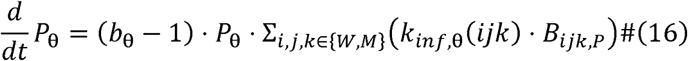

The initial conditions were:

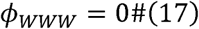

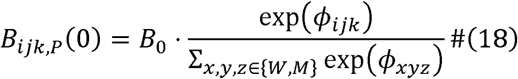

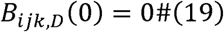

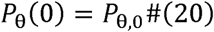

#### In vitro PKPD model parameter estimation

Monolix v2023R1 nonlinear mixed-effects modeling software (Lixoft, Antony, France) utilizing the SAEM algorithm was used to extend the model constructed above into a population model and estimate the parameter values. Variability between the 96-well plates and intra-plate variability were added to the parameters by assuming that each parameter followed a lognormal distribution. Residual variability was accounted for by using a combined proportional and additive model. The Monolix model file and the ‘.mlxtran’ file are provided in the supplementary files.

For the parameters *B*_0_, *k_growth_*, *k_death_*, *k_DP_*, *k_θ_*, *k_inf,θ,W_*, and *k_inf,θ,M_*, lognormal distributions were assumed for the individual parameters:

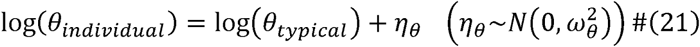

For the parameters log_10_ *C*, *ϕ*_WWW_*, ϕ*_WWM_*, ϕ*_WMW_*, ϕ*_WMM_*, ϕ*_MWW_, *ϕ*_MWM_, *ϕ*_MMW_, and *ϕ*_MMM_, a normal distribution was assumed for the individual parameters:

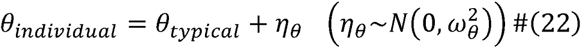

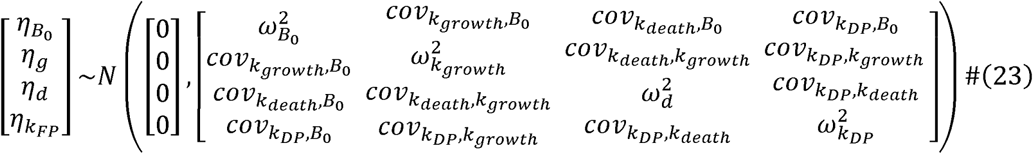

The residual variability was introduced using an additive error model:

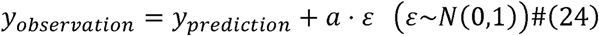

#### Biofilm Assays

##### Biofilm Formation

The ability of MP-A, PP-A, and the MP-A + PP-A cocktail to reduce biofilm formation was evaluated using a static biofilm assay. *Pseudomonas aeruginosa* (15-4) was grown in LB medium overnight at 37°C. The culture was diluted to attain an optical density at 600 nm (OD_600_) of 0.2 and transferred into 96-well polystyrene plates for biofilm formation. The plates were incubated at 37°C for 24 hours to allow biofilm establishment.

##### Phage Treatment of Biofilms

Following biofilm formation, the planktonic bacteria were gently removed, and the biofilm-containing wells were washed twice with phosphate-buffered saline (PBS) to remove non-adherent cells. Phages (MP-A, PP-A, or a combination of MP-A + PP-A) were then added at a total concentration of 2×10^10^ PFU/mL to the biofilm wells. Wells containing biofilms but not treated with phages served as controls. The plates were incubated for an additional 6 hours at 37°C.

##### Quantification of Biofilm Disruption

After phage treatment, the biofilm biomass was quantified using a crystal violet (CV) assay. Wells were washed three times with PBS to remove non-adherent cells, stained with 0.1% CV solution for 15 minutes, and then washed again with PBS to remove excess stain. The CV bound to the biofilm was solubilized using 30% acetic acid, and the OD was measured at 600 nm using the Spectramax ABS microplate reader (Molecular Devices, San Jose CA, USA).

Biofilm reduction was expressed as a percentage of the OD_600_ in the treated wells relative to the control wells. Each experiment was performed in duplicate, and results were averaged.

#### Accounting for Phage Saturation to Explain Results of Biofilm Assays

We explained the results of the biofilm assay by modifying the *in vitro* PKPD model. We assumed that the initial condition for the biofilm assay is the stationary state of bacteria. We simulated the *in vitro* PKPD model with the initial conditions (17), (18), and (19) in the absence of phages and considered the 24-hour dormant bacteria fraction, which was 0.9743, as the proportion of dormant bacteria in the biofilm.

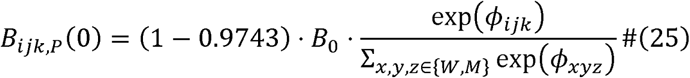

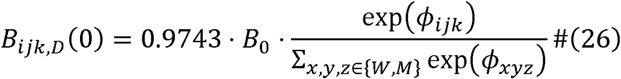

1A0ccording to the experimental protocol of the biofilm assay, we set B0 = 0.2 *OD*600 = 0.2 x 2 x 10^8^*CFU*/*mL*. For the regimens MP-A, PP-A, and MP-A + PP-A, we set (P1,0, P2,0)=(2 x 10^10^*PFU*/*mL*, 0*PFU*/*mL*), (0*PFU*/*mL*, 2 x 10^10^PFU/mL), and (10^10^*PFU*/*mL*, 10^10^PFU/mL)

Because changing the initial conditions was not sufficient to explain the differences in efficacy of the different regimens on the biofilm, we further modified the model to account for spatial heterogeneity using a saturation term 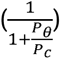 for the phage.

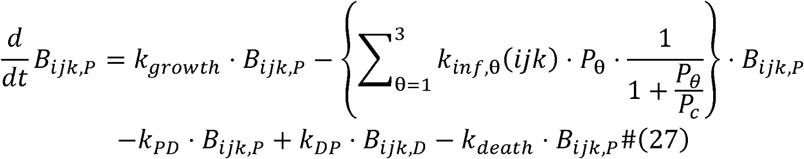

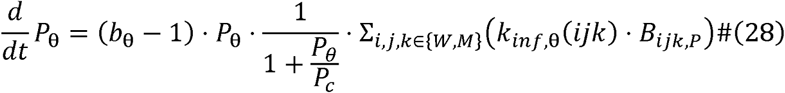

The phage concentration where the saturation effect is 1/2, which is .*P_c_*, was set to .*P_c_* = 1.5 x 10^7^ based on Roach et al.^25^.

#### In vivo mouse PKPD experiment

The PK of the MP-A + PP-A cocktail in the noninfected mouse model and the PKPD of the MP-A + PP-A cocktail in the infected mouse model were analyzed. The PKPD experiment in noninfected mice included a control group (three mice per timepoint) and three MP-A + PP-A treatment groups (three mice per timepoint), which were administered doses of 10^7^ PFU/head, 10^9^ PFU/head, and 10^11^ PFU/head, respectively. Similarly, the PKPD experiment in infected mice comprised a control group (four mice per timepoint) and three MP-A + PP-A treatment groups (four mice per timepoint), receiving doses of 10^7^ PFU/head, 10^7^ PFU/head, and 10^7^ PFU/head, respectively.

For the uninfected mouse model, the animals were fixed using a frame and using a 26-gauge needle, the control normal saline or the bacteriophage cocktail mixture was injected into the tail vein for one minute. The animals were sacrificed at 0, 0.5, 1, 4, 8, 12, 24, 48, and 60 hours after placebo or bacteriophage treatment.

For the infected mouse model, the bacterial mixture was prepared at a concentration of 10^7^ PFU/ 20 µL. The mice were fixed using the cervical skin fixation method, and the bacterial mixture was inoculated intranasally using a micropipette, 10 µL into each of the two nostrils. One hour after the infection, the animals were fixed using a frame and using a 26-gauge needle, the control normal saline or the bacteriophage cocktail mixture was injected into the tail vein for one minute. The animals were sacrificed at 0, 0.5, 1, 4, 8, 12, 24, 48, and 60 hours after placebo or bacteriophage treatment.

The animals to be sacrificed were anesthetized and the abdominal aorta was exposed using laparotomy. The blood was sampled from the abdominal aorta using a syringe and stored in a vacutainer tube with a clot activator. The blood sample was left for 15–20 minutes in room temperature for clotting. The clotted sample was centrifuged at 5000 rpm(7069 x g) for 5 minutes to extract the serum. After collection of serum, the animal was euthanized using CO_2_ and the lungs were extracted. The extracted lungs were measured for weight and were then homogenized using a bead homogenizer. The serum samples and homogenized lung samples were diluted and quantitated for bacterial load and phage titer.

#### Phage pharmacokinetics analysis

The initial concentration C_0_, area under the curve until the last timepoint AUC_last_, terminal half-life, initial concentration normalized by dose (C_0_/Dose), and area under the curve until the last timepoint normalized by dose (AUC_last_/Dose) were computed to characterize the pharmacokinetics of MP-A and PP-A in serum and lung tissue in both noninfected and infected mouse models. Phoenix® WinNonlin® software (Version 8.3, Certara, Radnor, PA, USA) was utilized for the calculation of these pharmacokinetic parameters, employing the linear trapezoidal method for AUC_last_ determination.

#### The in vivo PKPD model

The models developed and fitted to *in vitro* time-kill curves were extended to incorporate *in vivo* phage pharmacokinetics and immune system dynamics. The following system of ODE describes the updated model, where *θ* ∈ {1, 2, 3} and i,j,k ∈ {*W,M*}.

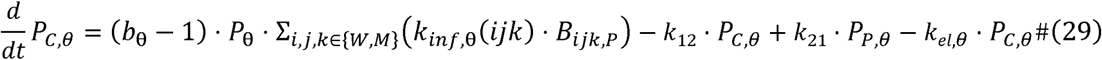

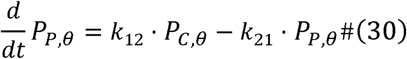

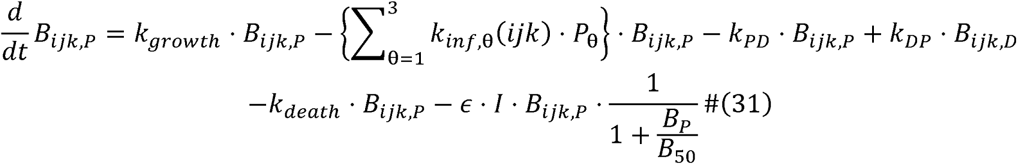

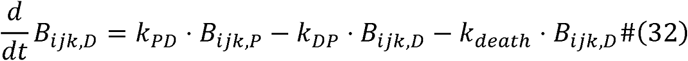

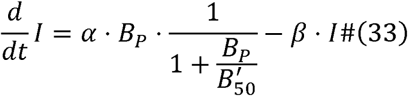

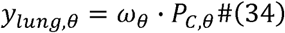

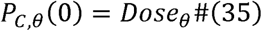

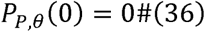

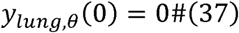

(*P_C,θ_*: Phage amount in the central compartment, *P_P,θ_*: Phage amount in the peripheral compartment; *I*: The immune sytem, *α* Induction rate constant for the immune system by total proliferative bacteria, *B*’_50_ : Load of proliferative bacteria needed to suppress the rate of immune system induction by half, *β*: Decay rate of the immune system, *ɛ*: Killing rate of proliferative bacteria by the immune system, *B*_50_: Load of proliferative bacteria needed to suppress the rate of immune system killing of bacteria by half, *y_lung,θ_*: Phage concentration in the lung compartment, *Dose_θ_*: The administered phage dose, *k*_12_: Distribution rate constant from central to peripheral compartment, *k*_21_: Distribution rate constant from peripheral to central compartment, *k_el,θ_*: phage elimination rate constant, *ω_θ_*: Proportionality factor linking *P_C,θ_*& to *y_lung,θ_*_)_

#### Parameter calibration of the mouse PKPD model

The calibration of the extra *in vivo* mouse PKPD parameters was conducted manually by using a grid search approach for each model parameter over a physiologically plausible range. Some of the parameters estimated from *in vitro* studies were also recalibrated to better describe the *in vivo* PKPD measurements. The recalibrated parameters included the proliferative capacity of the target bacteria, the infection rate of PP-A, and the burst size of PP-A. All other parameters were fixed to the values estimated from the *in vitro* data.

#### In vivo mouse lung bacterial load experiment

Two sets of experiments were conducted: one aimed at assessing the impact of phage type or cocktail composition and the other focused on evaluating the dose response of the MP-A + PP- A phage cocktail on lung bacterial load in intranasally infected mice treated with phages. For the first experiment, groups were organized as follows: a control group (n=6), a 5×10^9^ PFU/head MP-A treatment group (n=6), a 5×10^9^ PFU/head PP-A treatment group (n=6), and a 5×10^9^ PFU/head MP-A + PP-A treatment group (n=6). For the second experiment, groups were organized as follows: a control group (n=5), a 10^7^ PFU/head MP-A + PP-A treatment group (n=5), and a 10^11^ PFU/head MP-A + PP-A treatment group (n=5).

ICR mice were intranasally inoculated with 20 μL of prepared bacterial mixture. For the first experiment, bacterial doses of 6×10^7^ PFU/head were administered, and for the second experiment, doses of 1.5×10^8^ PFU/head were used. After a two-hour interval, either negative control saline or phage treatments were administered to the mice via the tail vein, with a volume of 200 μL. Upon the occurrence of the first mortality, all mice were euthanized, and lung tissues were collected. The time of first mortality was recorded as 21 hours for the first experiment and 20 hours for the second experiment. Bacterial quantification inside the lungs was performed using plating methods.

##### Assessing bacterial dose efficacy

Intranasal inoculation of mice with bacterial doses of 10^6^, 10^7^, 5×10^7^ CFU/head was performed to establish an acute pneumonia mouse model. Survival assessments were conducted at six-hour intervals. Each treatment group consisted of 10 replicates.

##### Evaluating the efficacy of the cocktail

A total of 5×10^8^ CFU/head of bacteria were inoculated intranasally into the mice to generate an acute pneumonia mouse model. For the control group, 0.2 mL of saline was injected into the tail vein and for the treatment group, 5×10^4^, 5×10^6^, 5×10^8^, and 5×10^10^ PFU/head of MP-A + PP-A cocktail was injected into the tail vein. Each control and treatment group had eight replicates.

Survival was evaluated at 0, 24, 48, 72, and 96 hours.

#### Stability Assays

##### Thermal Stability Assay

To assess thermal stability, phage suspensions of MP-A and PP-A were prepared at an initial concentration of 10^9^ PFU/mL. Phages were incubated at different temperatures (4°C, 25°C, 37°C, 50°C, 60°C, and 70°C) for 60 minutes. At the end of the incubation period, phage titers were determined using a standard double-layer agar method (plaque assay) against *Pseudomonas aeruginosa* host bacteria. For each temperature, three independent experiments were conducted. Phage viability was expressed as a percentage of the initial titer, and results were averaged across the replicates. Titer reductions were plotted as a function of temperature.

##### pH Stability Assay

The pH stability of MP-A and PP-A was tested by incubating phage suspensions (10^9^ PFU/mL) in buffer solutions of varying pH (pH 2, 3, 5, 6, 7, 8, 9, and 10). The phages were stored at 4°C for up to 30 days, and samples were collected at 1, 7, and 30 days for titer determination.

Phage titers were measured using plaque assays as described above, with the results averaged from three independent replicates. Phage viability was expressed as a percentage of the initial titer, and results were plotted to evaluate titer stability across the pH range over time.

### Simulations for strategies to overcome high dormant proportion for bacteria in the stationary phase

We simulated a more severe infection than that observed in the PKPD experiment by setting *B*_0_ = 0.04 *OD*600 = 0.04 x 2 10^8^ *CFU*/*mL*. We assumed that initial dormant bacteria do not exist and compared the case in which phage therapy with the MP-A + PP-A regimen with a total dose is administered immediately or 24 hours after the infection. When left untreated for 24 hours, most of the bacteria become dormant, and the efficacy of the delayed phage treatment dwindles. To explore phage-antibiotic combination therapy as a solution for this phenomenon, we added virtual antibiotic PKPD model components to our *in vivo* PKPD model.

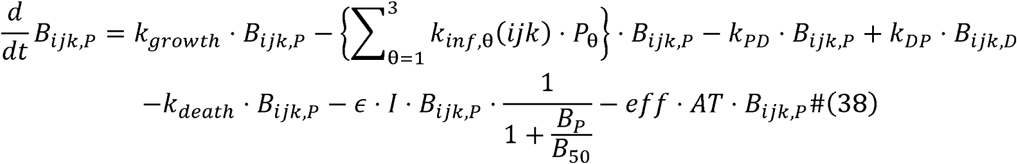

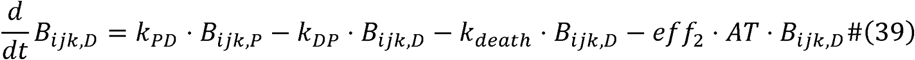

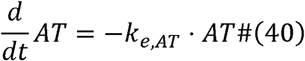

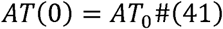

The antibiotic was assumed to have a killing effect proportional to the concentration of the antibiotic (AT) and the bacterial load, with different coefficients for proliferating bacteria and dormant bacteria (*eff* = 0.2 and *eff* = 0.1). The antibiotic was set to follow a 1-compartment distribution and linear elimination with elimination rate constant *k_e,AT_* = 1. The antibiotic was assumed to be administered via an intravenous bolus dose of *AT*_0_ = 10. We simulated the strategy of delayed phage treatment + single dose of antibiotic and delayed phage treatment + multiple doses of antibiotic with a 12-hour interval.

### QUANTIFICATION AND STATISTICAL ANALYSIS

The specifics of quantification and statistical analysis are delineated in the figure legends. Analysis of variance (ANOVA) and Tukey’s honestly significant difference (HSD) test were conducted using the Python package SciPy v1.11.2^48^. The log-rank test for survival was performed and analysis and Kaplan-Meier estimators for survival probability were computed using the Python package Lifelines v0.27.7^49^. Noncompartmental analysis for phage pharmacokinetics was executed using Phoenix® WinNonlin® v8.3 (Certara, PA, USA).

Nonlinear mixed-effects modeling of *in vitro* PKPD data was conducted using Monolix v2023R1 (Lixoft, Antony, France).

## Supplemental information titles and legends

Document S1. Figures S1–S8 and Tables S1–S4

