## Supplemental PDF for "Bridging *in vitro* and *in vivo* insights: A PKPD model for effective phage therapy against multidrug-resistant *Pseudomonas aeruginosa*"

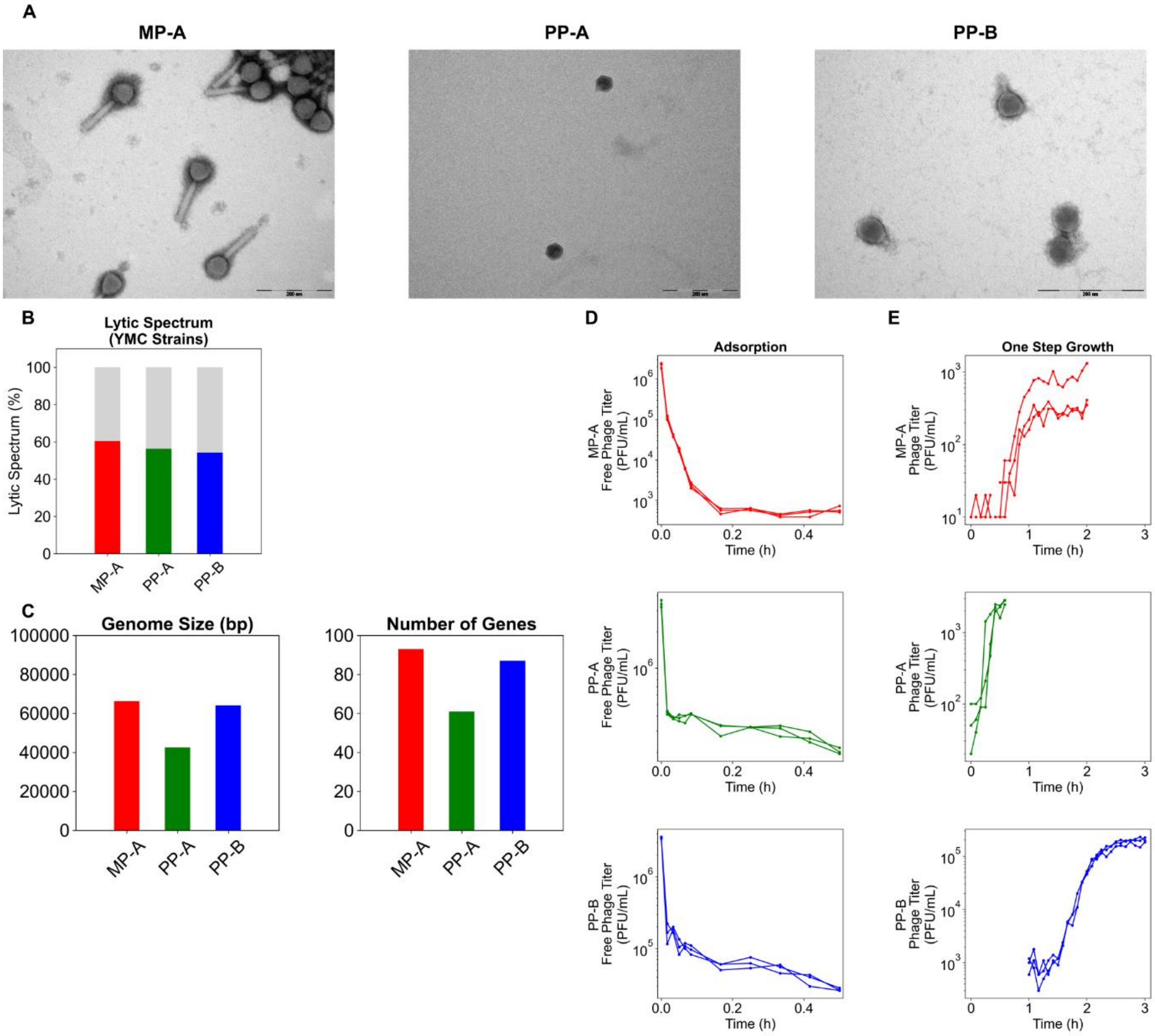


**Figure S1. Characterization of bacteriophages MP-A, PP-A, and PP-B.** (A) TEM images, (B) Lytic Spectrum, (C) genomics, and (D) adsorption and (E) one-step growth kinetics.

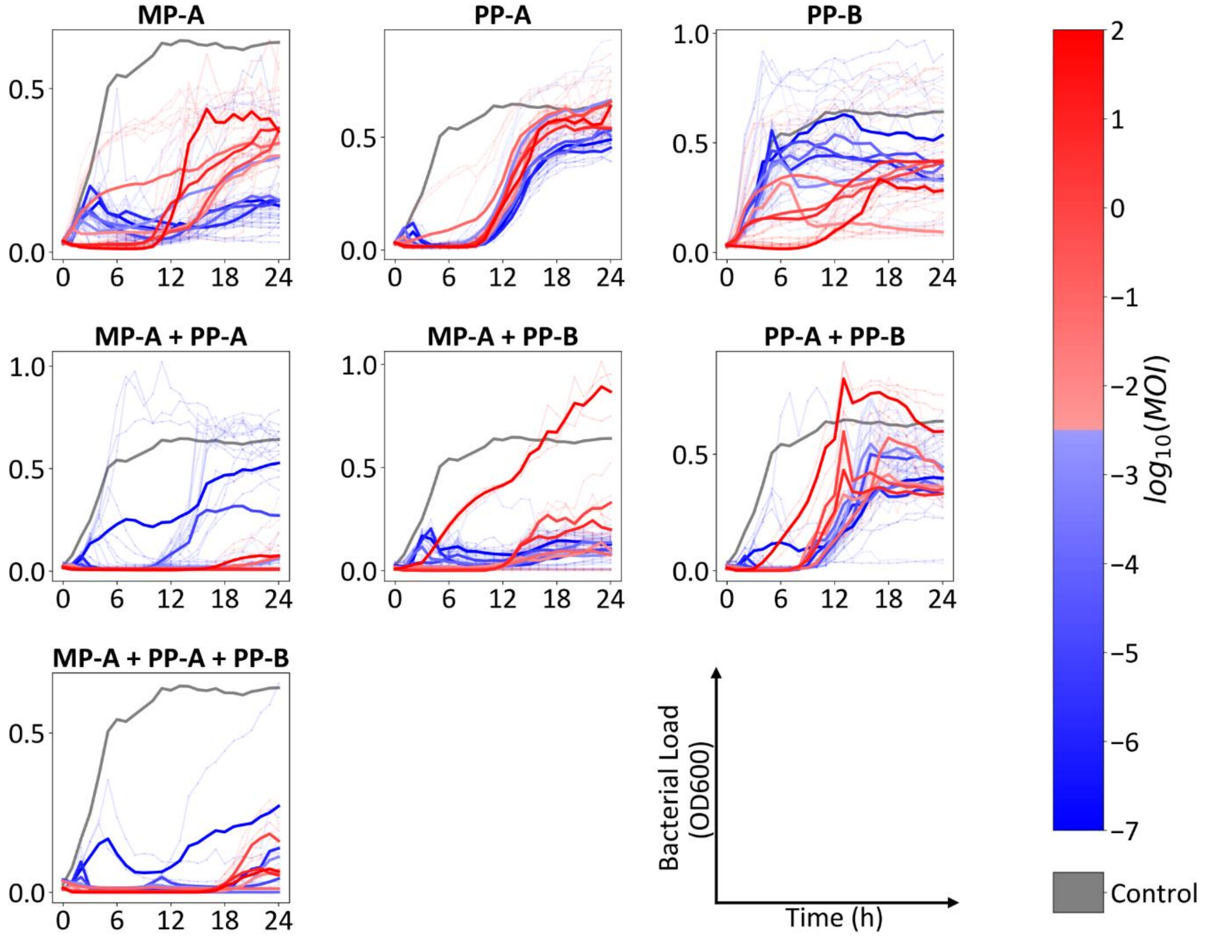


**Figure S2.** ***In vitro*** **efficacy of mono-phage and cocktail treatments**
*In vitro* time-kill assays were conducted for all treatments, with results presented using a color scale corresponding to the multiplicity of infection (MOI).

**Table S1.** Estimates of inter-plate variability in the model parameters

| **Parameter** | | **Unit** | **Estimate (RSE%)** |
|---|---|---|---|
| **Inter Plate Variability** | | | |
| $\omega_{B_0}$ | | **CV (%)** | **104.54 (3.73)** |
| $\omega_{k_{growth}}$ | | **CV (%)** | **32.06 (3.46)** |
| $\omega_{k_{death}}$ | | **CV (%)** | **207.38 (5.44)** |
| $\omega_{k_{DP}}$ | | **CV (%)** | **255.25 (4.92)** |
| $\omega_{log_{10}(C)}$ | | **CV (%)** | **3.25 (4.59)** |
| $\omega_{log_{10}(\phi_{WWM})}$ | | **CV (%)** | **48.33 (5.49)** |
| $\omega_{log_{10}(\phi_{WMW})}$ | | **CV (%)** | **24.53 (7.55)** |
| $\omega_{log_{10}(\phi_{WMM})}$ | | **CV (%)** | **32.15 (6.63)** |
| $\omega_{log_{10}(\phi_{MWW})}$ | | **CV (%)** | **16.05 (10.0)** |
| $\omega_{log_{10}(\phi_{MWM})}$ | | **CV (%)** | **45.29 (5.44)** |
| $\omega_{log_{10}(\phi_{MMW})}$ | | **CV (%)** | **32.67 (7.64)** |
| $\omega_{log_{10}(\phi_{MMM})}$ | | **CV (%)** | **36.33 (6.68)** |
| $\omega_{log_{10}(k_{inf,W})}$ | **MP-A** | **CV (%)** | **1.72 (13.8)** |
| | **PP-A** | **CV (%)** | **3.03 (10.4)** |
| | **PP-B** | **CV (%)** | **4.38 (9.68)** |
| $\omega_{log_{10}(k_{inf,M})}$ | **MP-A** | **CV (%)** | **0.96 (33.6)** |
| | **PP-A** | **CV (%)** | **3.12 (35.2)** |
| | **PP-B** | **CV (%)** | **4.53 (15.3)** |
| $\omega_b$ | **MP-A** | **CV (%)** | **52.22 (7.56)** |
| | **PP-A** | **CV (%)** | **35.42 (10.7)** |
| | **PP-B** | **CV (%)** | **112.3 (8.22)** |
| **Correlations** | | | |
| $\rho_{k_{death},B_0}$ | | | **-0.23 (28.4)** |
| $\rho_{k_{growth},B_0}$ | | | **-0.35 (12.9)** |
| $\rho_{k_{DP},B_0}$ | | | **-0.054 (118)** |
| $\rho_{log_{10}(C),B_0}$ | | | **0.22 (26.3)** |
| $\rho_{k_{growth},k_{death}}$ | | | **0.33 (17.8)** |
| $\rho_{k_{DP},k_{death}}$ | | | **-0.2 (40.0)** |
| $\rho_{log_{10}(C),k_{death}}$ | | | **0.43 (14.7)** |
| $\rho_{k_{DP},k_{growth}}$ | | | **0.1 (59.1)** |
| $\rho_{log_{10}(C),k_{growth}}$ | | | **-0.34 (15.2)** |
| $\rho_{log_{10}(C),k_{DP}}$ | | | **-0.67 (6.33)** |
| **Residual Variability** | | | |
| $a$ | | | **0.026 (0.753)** |

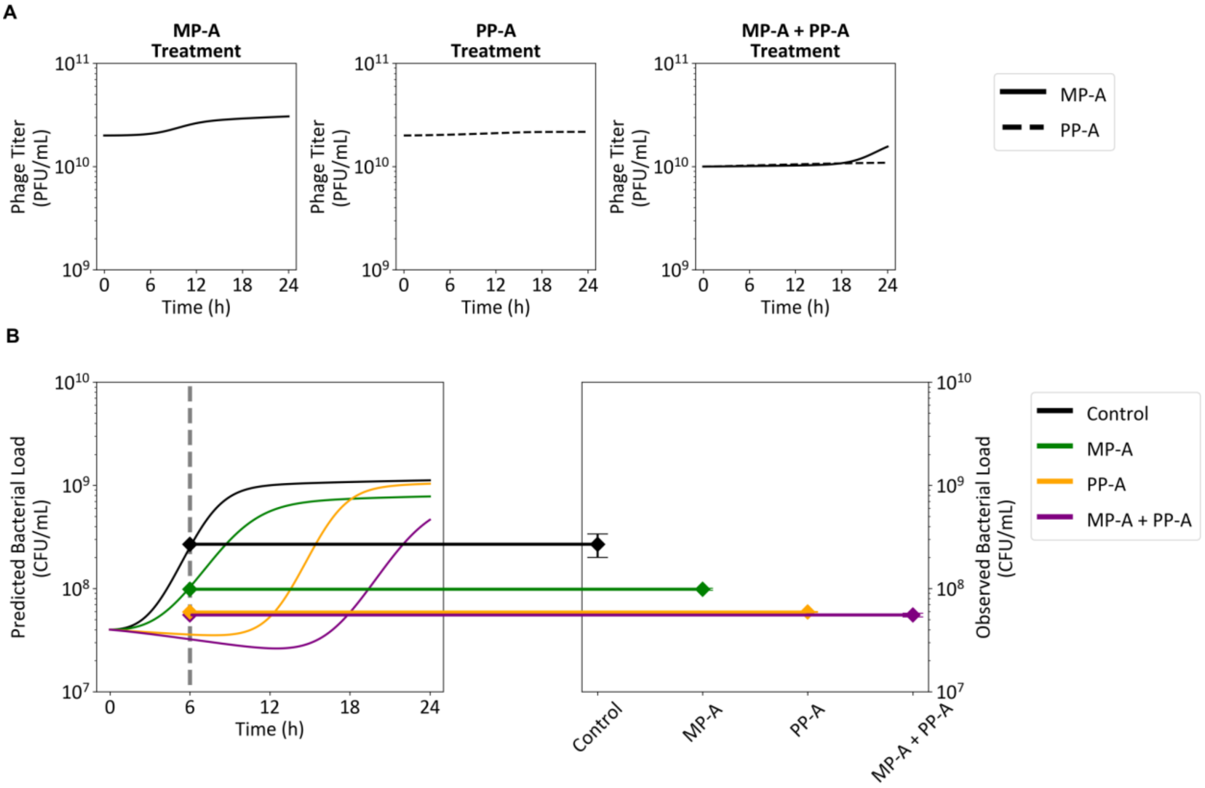


**Figure S3. Mathematical modeling of biofilm elimination**
The *in vitro* model was modified to account for differences in efficacy between planktonic and biofilm bacteria and among phage regimens. (A) The predicted phage titers for each regimen are shown. (B) Predicted bacterial load (left panel) and observed bacterial load (right panel) are compared. The observed data (right panel) are shown using error bars with average±standard deviation of the bacterial load. The number of samples for each treatment group was n=4.

**Table S2.** Noncompartmental analysis of healthy mice PK

(A) MP-A, Serum

| | G2, $10^7$ PFU/head | G3, $10^9$ PFU/head | G4, $10^{11}$ PFU/head |
|---|---|---|---|
| $C_0$ (PFU/mL) | $1.53 \times 10^2$ | $7.86 \times 10^6$ | $5.61 \times 10^9$ |
| $AUC_{last}$ (PFU h/mL) | $3.72 \times 10^2$ | $3.43 \times 10^6$ | $2.47 \times 10^9$ |
| Terminal half-life (h) | 5.2 | 5.6 | 2.5 |
| $C_0/Dose$ (/mL) | $1.53 \times 10^{-5}$ | $7.86 \times 10^{-3}$ | $5.61 \times 10^{-2}$ |
| $AUC_{last}/Dose$ (h/mL) | $3.72 \times 10^{-5}$ | $3.43 \times 10^{-3}$ | $2.47 \times 10^{-2}$ |

(B) MP-A, Lung

| | G2, $10^7$ PFU/head | G3, $10^9$ PFU/head | G4, $10^{11}$ PFU/head |
|---|---|---|---|
| $C_0$ (PFU/mL) | $2.50 \times 10^1$ | $2.03 \times 10^5$ | $9.63 \times 10^8$ |
| $AUC_{last}$ (PFU h/mL) | $1.21 \times 10^3$ | $1.69 \times 10^6$ | $3.11 \times 10^9$ |
| Terminal half-life (h) | 25.6 | 17.5 | 10.4 |
| $C_0/Dose$ (/mL) | $2.50 \times 10^{-6}$ | $2.03 \times 10^{-4}$ | $9.63 \times 10^{-3}$ |
| $AUC_{last}/Dose$ (h/mL) | $1.21 \times 10^{-4}$ | $1.69 \times 10^{-3}$ | $3.11 \times 10^{-2}$ |
| Lung/Serum $C_0$ ratio | 0.16 | 0.026 | 0.17 |

(C) PP-A, Serum

| | G2, $10^7$ PFU/head | G3, $10^9$ PFU/head | G4, $10^{11}$ PFU/head |
|---|---|---|---|
| $C_0$ (PFU/mL) | $2.84 \times 10^7$ | $5.74 \times 10^9$ | $6.38 \times 10^{11}$ |
| $AUC_{last}$ (PFU h/mL) | $2.98 \times 10^7$ | $2.84 \times 10^9$ | $4.63 \times 10^{11}$ |
| Terminal half-life (h) | 1.1 | 1.8 | 1.8 |
| $C_0/Dose$ (/mL) | $2.84 \times 10^0$ | $5.74 \times 10^0$ | $6.38 \times 10^0$ |
| $AUC_{last}/Dose$ (h/mL) | $2.98 \times 10^0$ | $2.84 \times 10^0$ | $4.63 \times 10^0$ |

(D) PP-A, Lung

| | G2, $10^7$ PFU/head | G3, $10^9$ PFU/head | G4, $10^{11}$ PFU/head |
|---|---|---|---|
| $C_0$ (PFU/mL) | $4.21 \times 10^5$ | $4.78 \times 10^7$ | $5.13 \times 10^9$ |
| $AUC_{last}$ (PFU h/mL) | $5.78 \times 10^5$ | $4.82 \times 10^7$ | $1.50 \times 10^{10}$ |
| Terminal half life (h) | 8.7 | 18.1 | 6.9 |
| $C_0/Dose$ (/mL) | $4.21 \times 10^{-2}$ | $4.78 \times 10^{-2}$ | $5.13 \times 10^{-2}$ |
| $AUC_{last}/Dose$ (h/mL) | $5.78 \times 10^{-2}$ | $4.82 \times 10^{-2}$ | $1.50 \times 10^{-1}$ |
| Lung/Serum $C_0$ ratio | 0.015 | 0.0083 | 0.0080 |

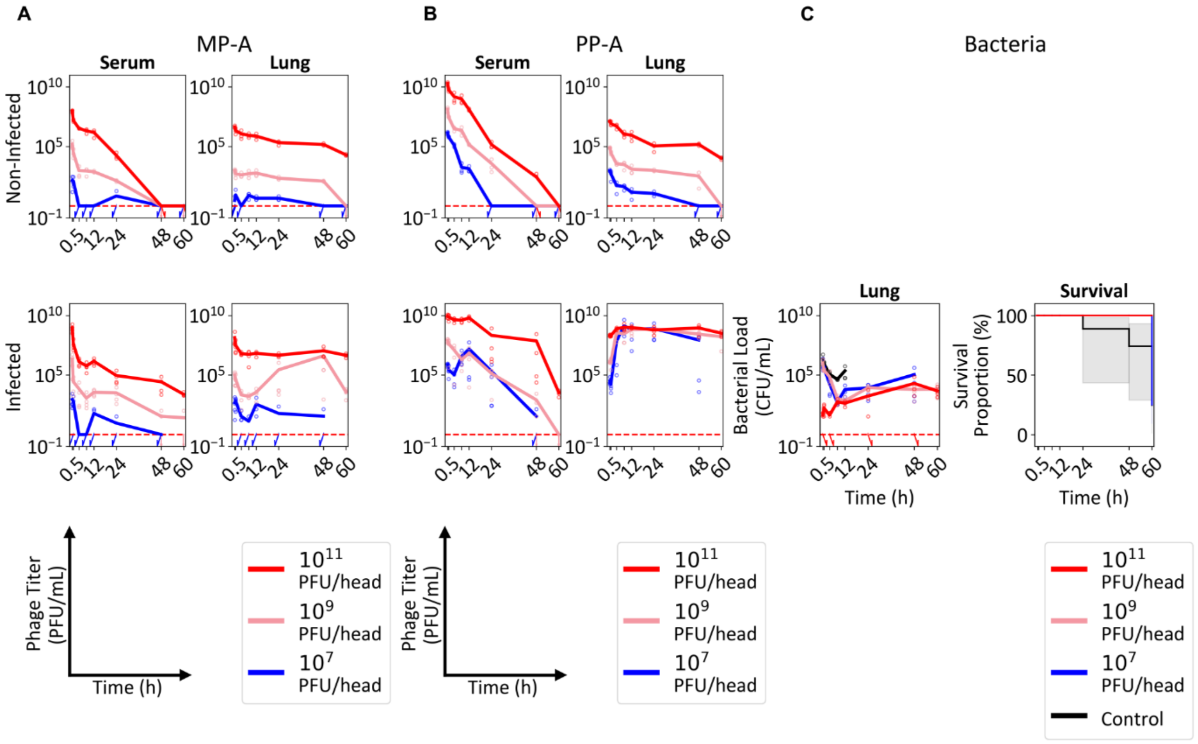


**Figure S4. *In vivo* PK/PD and survival**
PKPD profiles were evaluated in uninfected mice and mice with acute pneumonia. The time-course dynamics of MP-A (A), PP-A (B), and bacteria (C) are illustrated. Survival outcomes are additionally depicted using a Kaplan-Meier plot. Measurements below the Lower Limit of Quantification (LLOQ) (phage: 1 PFU/mL, bacteria: 1 CFU/mL) were imputed with the LLOQ value when calculating the mean for each respective timepoint. The meas for each timepoint are connected with solid lines.

**Table S3.** Noncompartmental analysis of infected mice PK

(A) MP-A, Serum

| | G2, $10^7$ PFU/head | G3, $10^9$ PFU/head | G4, $10^{11}$ PFU/head |
|---|---|---|---|
| $C_0$ (PFU/mL) | $1.41 \times 10^3$ | $1.87 \times 10^9$ | $2.18 \times 10^{11}$ |
| $AUC_{last}$ (PFU h/mL) | $2.62 \times 10^3$ | $4.81 \times 10^8$ | $6.06 \times 10^{10}$ |
| Terminal half life (h) | 3.7 | 5.8 | 6.5 |
| $C_0/Dose$ (/mL) | $1.41 \times 10^{-4}$ | $1.87 \times 10^0$ | $2.18 \times 10^0$ |
| $AUC_{last}/Dose$ (h/mL) | $2.62 \times 10^{-4}$ | $4.81 \times 10^{-1}$ | $6.06 \times 10^{-1}$ |
| Infected/Noninfected $AUC_{last}$ | 7.04 | 140.23 | 24.53 |

(B) MP-A, Lung

| | G2, $10^7$ PFU/head | G3, $10^9$ PFU/head | G4, $10^{11}$ PFU/head |
|---|---|---|---|
| $C_0$ (PFU/mL) | $5.82 \times 10^2$ | $2.16 \times 10^6$ | $6.63 \times 10^9$ |
| $AUC_{last}$ (PFU h/mL) | $8.45 \times 10^7$ | $8.74 \times 10^8$ | $7.46 \times 10^9$ |
| Terminal half life (h) | NA | NA | 28.8 |
| $C_0/Dose$ (/mL) | $5.82 \times 10^{-5}$ | $2.16 \times 10^{-3}$ | $6.63 \times 10^{-2}$ |
| $AUC_{last}/Dose$ (h/mL) | $8.45 \times 10^0$ | $8.74 \times 10^{-1}$ | $7.46 \times 10^{-2}$ |
| Lung/Serum $C_0$ ratio | 0.37 | 0.0012 | 0.03 |
| Infected/Noninfected $AUC_{last}$ | $6.98 \times 10^5$ | 517.16 | 2.40 |

(C) PP-A, Serum

| | G2, $10^7$ PFU/head | G3, $10^9$ PFU/head | G4, $10^{11}$ PFU/head |
|---|---|---|---|
| $C_0$ (PFU/mL) | $4.79 \times 10^7$ | $3.34 \times 10^9$ | $2.40 \times 10^{11}$ |
| $AUC_{last}$ (PFU h/mL) | $3.59 \times 10^9$ | $6.40 \times 10^9$ | $2.66 \times 10^{12}$ |
| Terminal half life (h) | 1.9 | 2.8 | 3.5 |
| $C_0/Dose$ (/mL) | $4.79 \times 10^0$ | $3.34 \times 10^0$ | $2.40 \times 10^0$ |
| $AUC_{last}/Dose$ (h/mL) | $3.59 \times 10^2$ | $6.40 \times 10^0$ | $2.66 \times 10^1$ |
| Infected/Noninfected $AUC_{last}$ | 120.47 | 2.25 | 5.75 |

(D) PP-A, Lung

| | G2, $10^7$ PFU/head | G3, $10^9$ PFU/head | G4, $10^{11}$ PFU/head |
|---|---|---|---|
| $C_0$ (PFU/mL) | $1.62 \times 10^6$ | $3.17 \times 10^7$ | $5.53 \times 10^9$ |
| $AUC_{last}$ (PFU h/mL) | $4.02 \times 10^{12}$ | $6.68 \times 10^{11}$ | $1.10 \times 10^{12}$ |
| Terminal half life (h) | NA | 16.8 | 44.6 |
| $C_0/Dose$ (/mL) | $1.62 \times 10^{-1}$ | $3.17 \times 10^{-2}$ | $5.53 \times 10^{-2}$ |
| $AUC_{last}/Dose$ (h/mL) | $4.02 \times 10^5$ | $6.68 \times 10^2$ | $1.10 \times 10^1$ |
| Lung/Serum $C_0$ ratio | 0.034 | 0.0095 | 0.023 |
| Infected/Noninfected $AUC_{last}$ | $6.96 \times 10^6$ | $1.39 \times 10^4$ | 73.33 |

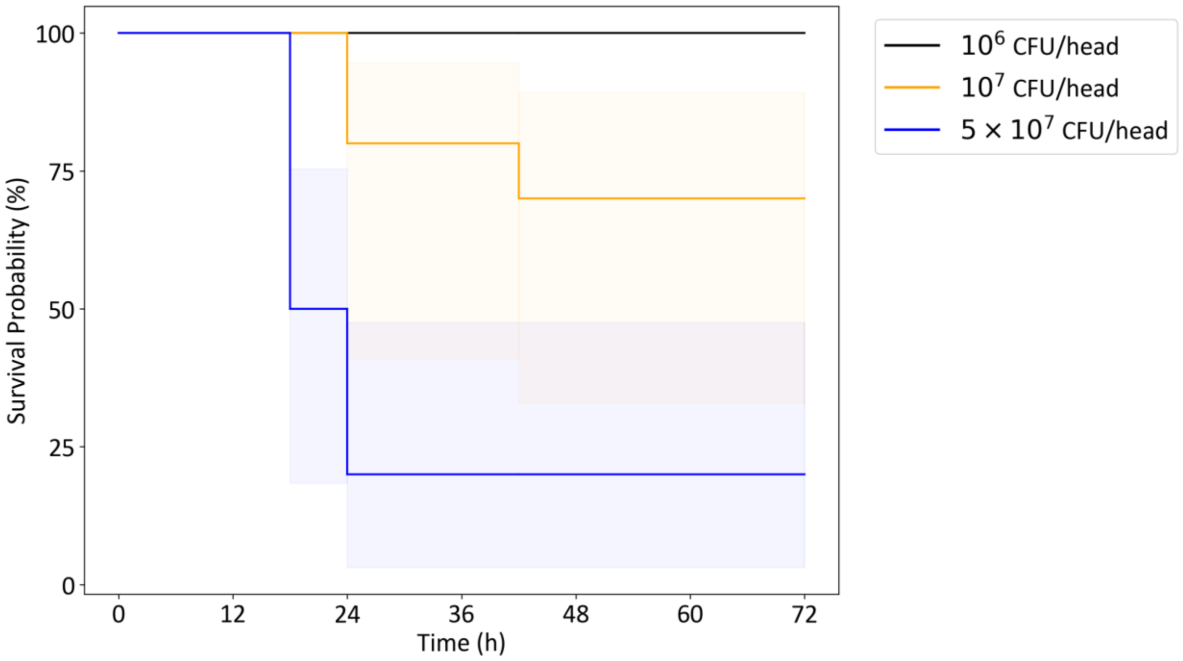


**Figure S5 Effect of *P. aeruginosa* dose on survival in acute pneumonia model**
The Kaplan-Meier plot depicts the survival of mice in relation to varying levels of bacterial load.

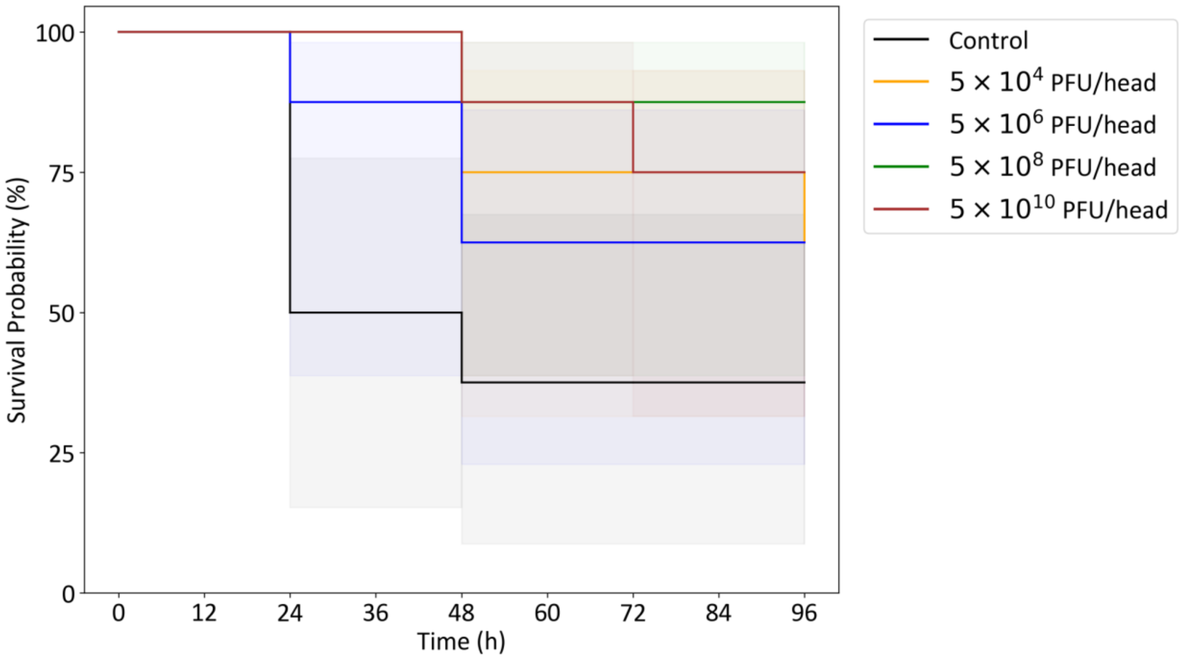


**Figure S6. Effect of MP-A+PP-A cocktail dose on survival of acute pneumonia mouse model**
The Kaplan-Meier plot illustrates the survival of mice based on the dosage of administered phage.

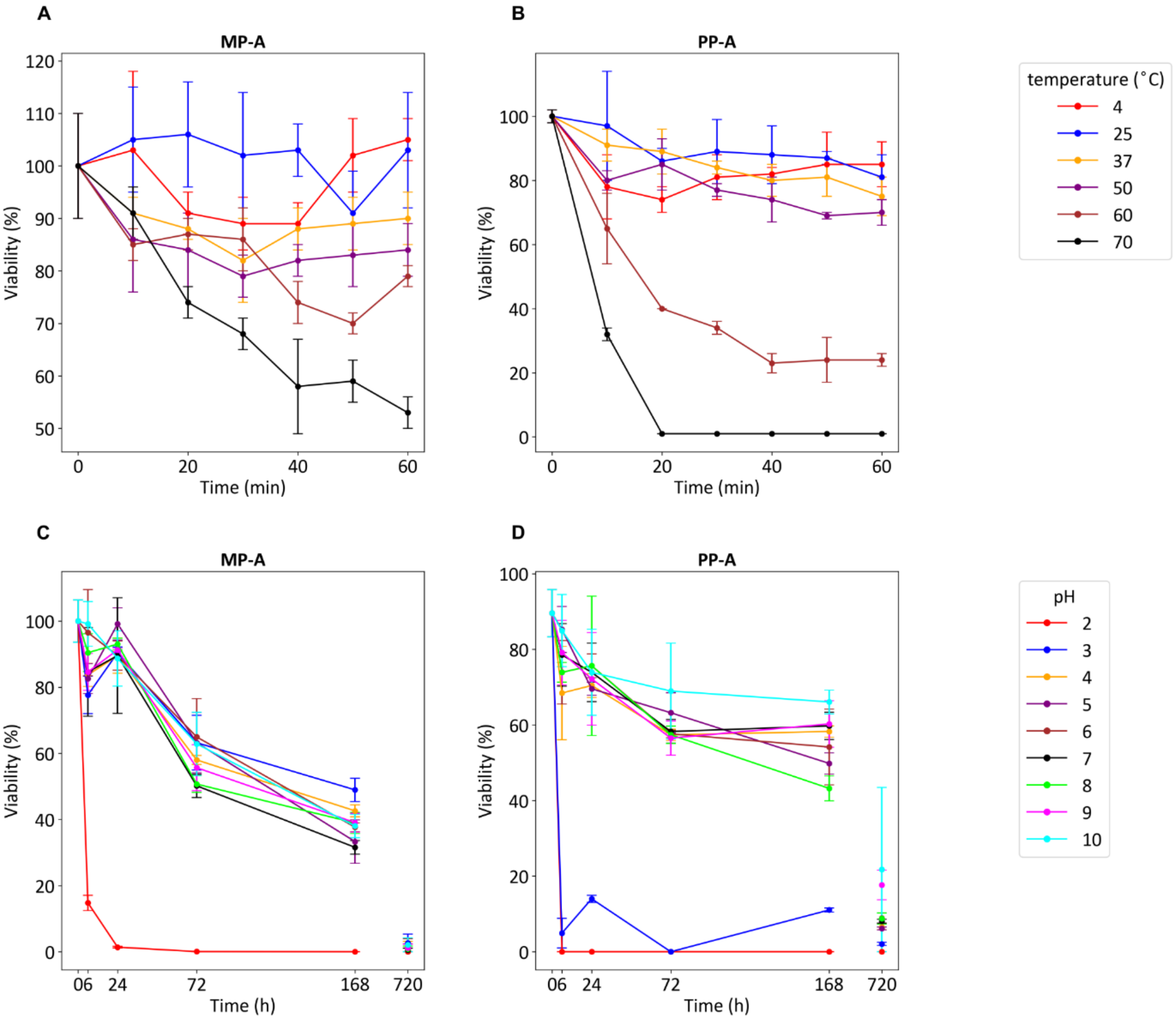


**Figure S7. Phage thermal and pH stability**
The stability of the constituent phages of the MP-A + PP-A cocktail was evaluated. The thermal stability of the strains MP-A (A) and PP-A (B) are shown. The pH stability of the strains MP-A (C) and PP-A (D) are shown. Error bars represent average±standard deviation of the viability proportion.

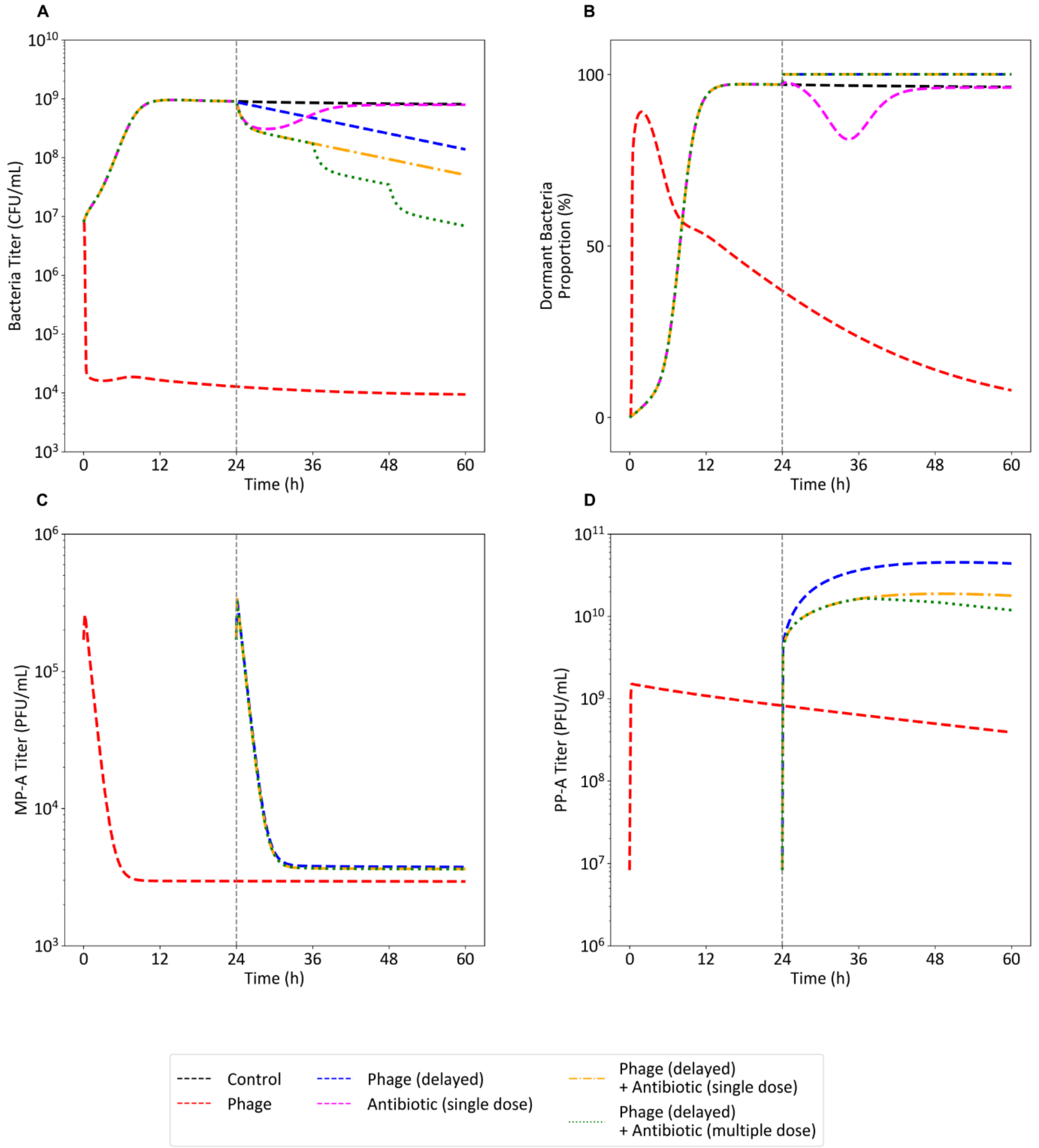


**Figure S8. Simulation of strategies to overcome high dormant fraction of bacteria at the stationary phase.**
Phage administration when the bacteria is in the stationary phase (blue) is ineffective compared to phage administration when bacteria has not reached the stationary phase (red). This inefficacy is due to the high dormant fraction of bacteria in the stationary phase. Strategies to overcome this inefficacy such as combination therapy with a single dose (orange) or multiple doses (green) of antibiotic are explored through simulations. Simulated bacterial load (A), Dormant fraction (B), MP-A titer (C), and PP-A titer (D) are shown.
.

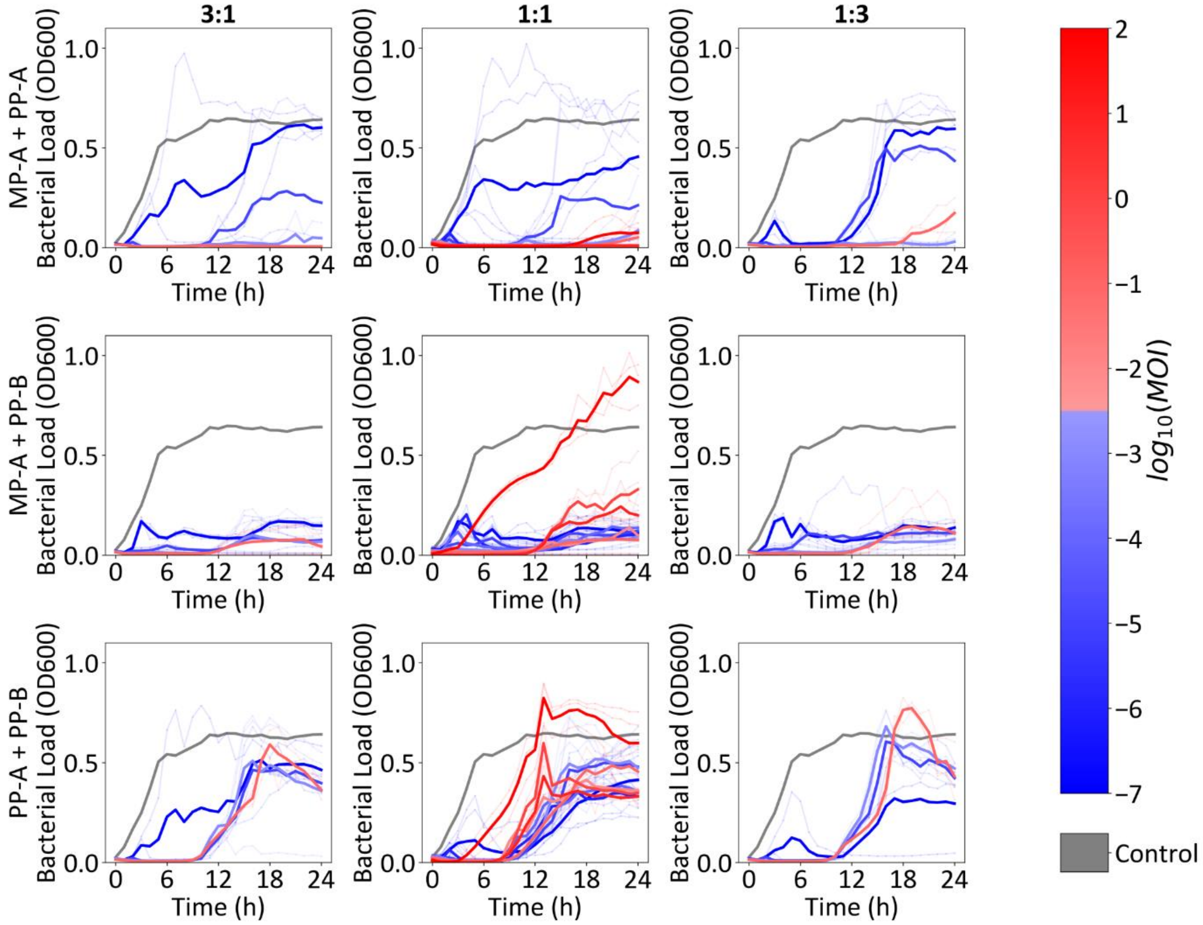


**Figure S9. Time-kill assay results for double cocktails stratified by cocktail components and mixing ratio**

*In vitro* time-kill assay results for phage double cocktails were stratified by cocktail composition and mixing ratio, with results presented using a color scale corresponding to the multiplicity of infection (MOI).

**Table S4 Experimental Setup for Time-Kill Assay Protocol1**

| 96well Plate Number | Treatment Group | | | Number of Replicates |
|---|---|---|---|---|
| | Treatment Type | MOI | RATIO | |
| 1 | Control | 0 | - | 3 |
| | MP-A + PP-B | $10^{-5}$ | 3:1 | 3 |
| | | | 1:1 | 3 |
| | | | 1:3 | 3 |
| | | $10^{-3}$ | 3:1 | 3 |
| | | | 1:1 | 3 |
| | | | 1:3 | 3 |
| | | $10^{-1}$ | 3:1 | 3 |
| | | | 1:1 | 3 |
| | | | 1:3 | 3 |
| 2 | Control | 0 | - | 3 |
| | MP-A + PP-B | $10^{-7}$ | 3:1 | 3 |
| | | | 1:1 | 3 |
| | | | 1:3 | 3 |
| | | $10^{-5}$ | 3:1 | 3 |
| | | | 1:1 | 3 |
| | | | 1:3 | 3 |
| | | $10^{-3}$ | 3:1 | 3 |
| | | | 1:1 | 3 |
| | | | 1:3 | 3 |
| | | $10^{-1}$ | 3:1 | 3 |
| | | | 1:1 | 3 |
| | | | 1:3 | 3 |
| | Control | 0 | - | 3 |
| 3 | PP-A + PP-B | $10^{-7}$ | 3:1 | 3 |
| | | | 1:1 | 3 |
| | | | 1:3 | 3 |
| | | $10^{-5}$ | 3:1 | 3 |
| | | | 1:1 | 3 |
| | | | 1:3 | 3 |
| | | $10^{-3}$ | 3:1 | 3 |
| | | | 1:1 | 3 |
| | | | 1:3 | 3 |
| | | $10^{-1}$ | 3:1 | 3 |
| | | | 1:1 | 3 |
| | | | 1:3 | 3 |
| 4 | Control | 0 | - | 3 |
| | MP-A + PP-A | $10^{-7}$ | 3:1 | 3 |
| | | | 1:1 | 3 |
| | | | 1:3 | 3 |
| | | $10^{-5}$ | 3:1 | 3 |
| | | | 1:1 | 3 |
| | | | 1:3 | 3 |
| | | $10^{-3}$ | 3:1 | 3 |
| | | | 1:1 | 3 |
| | | | 1:3 | 3 |
| | | $10^{-1}$ | 3:1 | 3 |
| | | | 1:1 | 3 |
| | | | 1:3 | 3 |
| 5 | Control | 0 | - | 3 |
| | PP-B | $10^{-7}$ | - | 3 |
| | | $10^{-5}$ | - | 3 |
| | | $10^{-3}$ | - | 3 |

| | | | | |
|---|---|---|---|---|
| | | $10^{-1}$ | - | 3 |
| | MP-A+PP-B | $10^{-7}$ | 1:1 | 3 |
| | | $10^{-5}$ | 1:1 | 3 |
| | | $10^{-3}$ | 1:1 | 3 |
| | | $10^{-1}$ | 1:1 | 3 |
| 6 | Control | 0 | - | 3 |
| | MP-A | $10^{-7}$ | - | 3 |
| | | $10^{-5}$ | - | 3 |
| | | $10^{-3}$ | - | 3 |
| | | $10^{-1}$ | - | 3 |
| | PP-A + PP-B | $10^{-7}$ | 1:1 | 3 |
| | | $10^{-5}$ | 1:1 | 3 |
| | | $10^{-3}$ | 1:1 | 3 |
| | | $10^{-1}$ | 1:1 | 3 |
| 7 | Control | 0 | - | 3 |
| | MP-A | $10^{-7}$ | - | 3 |
| | | $10^{-5}$ | - | 3 |
| | | $10^{-3}$ | - | 3 |
| | | $10^{-1}$ | - | 3 |
| | MP-A + PP-A | $10^{-7}$ | 1:1 | 3 |
| | | $10^{-5}$ | 1:1 | 3 |
| | | $10^{-3}$ | 1:1 | 3 |
| | | $10^{-1}$ | 1:1 | 3 |
| 8 | Control | 0 | - | 3 |
| | PP-A | $10^{-7}$ | - | 3 |
| | | $10^{-5}$ | - | 3 |
| | | $10^{-3}$ | - | 3 |
| | | $10^{-1}$ | - | 3 |
| | MP-A + PP-A + PP-B | $10^{-7}$ | 1:1:1 | 3 |
| | | $10^{-5}$ | 1:1:1 | 3 |
| | | $10^{-3}$ | 1:1:1 | 3 |
| | | $10^{-1}$ | 1:1:1 | 3 |
| 9 | Control | 0 | - | 6 |
| | PP-B | $10^{-7}$ | - | 3 |
| | | $10^{-5}$ | - | 3 |
| | | $10^{-3}$ | - | 3 |
| | | $10^{-1}$ | - | 3 |
| | | $10^{0}$ | - | 3 |
| | | $10^{1}$ | - | 3 |
| | MP-A | $10^{-7}$ | - | 3 |
| | | $10^{-5}$ | - | 3 |
| | | $10^{-3}$ | - | 3 |
| | | $10^{-1}$ | - | 3 |
| | | $10^{0}$ | - | 3 |
| | | $10^{1}$ | - | 3 |
| | PP-A | $10^{-7}$ | - | 3 |
| | | $10^{-5}$ | - | 3 |
| | | $10^{-3}$ | - | 3 |
| | | $10^{-1}$ | - | 3 |
| | | $10^{0}$ | - | 3 |
| | | $10^{1}$ | - | 3 |
| 10 | Control | 0 | - | 6 |
| | PP-B | $10^{-7}$ | - | 3 |
| | | $10^{-5}$ | - | 3 |

| | | | | |
|---|---|---|---|---|
| | | $10^{-3}$ | - | 3 |
| | | $10^{-1}$ | - | 3 |
| | | $10^{0}$ | - | 3 |
| | | $10^{1}$ | - | 3 |
| | MP-A | $10^{-7}$ | - | 3 |
| | | $10^{-5}$ | - | 3 |
| | | $10^{-3}$ | - | 3 |
| | | $10^{-1}$ | - | 3 |
| | | $10^{0}$ | - | 3 |
| | | $10^{1}$ | - | 3 |
| | PP-A | $10^{-7}$ | - | 3 |
| | | $10^{-5}$ | - | 3 |
| | | $10^{-3}$ | - | 3 |
| | | $10^{-1}$ | - | 3 |
| | | $10^{0}$ | - | 3 |
| | | $10^{1}$ | - | 3 |
| 11 | Control | 0 | - | 3 |
| | MP-A + PP-A | $10^{-6}$ | 1:1 | 3 |
| | | $10^{-4}$ | 1:1 | 3 |
| | | $10^{-2}$ | 1:1 | 3 |
| | | $10^{0}$ | 1:1 | 3 |
| | | $10^{1}$ | 1:1 | 3 |
| | | $10^{2}$ | 1:1 | 3 |
| 12 | Control | 0 | - | 3 |
| | PP-B | $10^{-6}$ | - | 3 |
| | | $10^{-4}$ | - | 3 |
| | | $10^{-2}$ | - | 3 |
| | | $10^{0}$ | - | 3 |
| | | $10^{1}$ | - | 3 |
| | | $10^{2}$ | - | 3 |
| | MP-A | $10^{-6}$ | - | 3 |
| | | $10^{-4}$ | - | 3 |
| | | $10^{-2}$ | - | 3 |
| | | $10^{0}$ | - | 3 |
| | | $10^{1}$ | - | 3 |
| | | $10^{2}$ | - | 3 |
| | PP-A | $10^{-6}$ | - | 3 |
| | | $10^{-4}$ | - | 3 |
| | | $10^{-2}$ | - | 3 |
| | | $10^{0}$ | - | 3 |
| | | $10^{1}$ | - | 3 |
| | | $10^{2}$ | - | 3 |
| 13 | Control | 0 | - | 3 |
| | MP-A + PP-A + PP-B | $10^{-6}$ | 1:1:1 | 3 |
| | | $10^{-4}$ | 1:1:1 | 3 |
| | | $10^{-2}$ | 1:1:1 | 3 |
| | | $10^{0}$ | 1:1:1 | 3 |
| | | $10^{1}$ | 1:1:1 | 3 |
| | | $10^{2}$ | 1:1:1 | 3 |
| 14 | Control | 0 | - | 3 |
| | MP-A + PP-B | $10^{-6}$ | 1:1 | 3 |
| | | $10^{-4}$ | 1:1 | 3 |
| | | $10^{-2}$ | 1:1 | 3 |
| | | $10^{0}$ | 1:1 | 3 |

| | | | | |
|---|---|---|---|---|
| | | $10^{1}$ | 1:1 | 3 |
| | | $10^{2}$ | 1:1 | 3 |
| | PP-A + PP-B | $10^{-6}$ | 1:1 | 3 |
| | | $10^{-4}$ | 1:1 | 3 |
| | | $10^{-2}$ | 1:1 | 3 |
| | | $10^{0}$ | 1:1 | 3 |
| | | $10^{1}$ | 1:1 | 3 |
| | | $10^{2}$ | 1:1 | 3 |
